# Structural basis for the inhibition of the *Bacillus subtilis* c-di-AMP cyclase CdaA by the phosphoglucomutase GlmM

**DOI:** 10.1101/2021.05.26.445777

**Authors:** Monisha Pathania, Tommaso Tosi, Charlotte Millership, Fumiya Hoshiga, Rhodri M. L. Morgan, Paul S. Freemont, Angelika Gründling

## Abstract

Cyclic-di-adenosine monophosphate (c-di-AMP) is an important nucleotide signalling molecule, which plays a key role in osmotic regulation in bacteria. Cellular c-di-AMP levels are tightly regulated, as both high and low levels have a negative impact on bacterial growth. Here, we investigated how the activity of the main *Bacillus subtilis* c-di-AMP cyclase CdaA is regulated by the phosphoglucomutase GlmM. c-di-AMP is produced from two molecules of ATP by proteins containing a deadenylate cyclase (DAC) domain. CdaA is a membrane-linked cyclase with an N-terminal transmembrane domain followed by the cytoplasmic DAC domain. Here we show, using the soluble catalytic *B. subtilis* CdaA_CD_ domain and purified full-length GlmM or the GlmM_F369_ variant lacking the C-terminal flexible domain 4, that the cyclase and phosphoglucomutase form a stable complex *in vitro* and that GlmM is a potent cyclase inhibitor. We determined the crystal structure of the individual *B. subtilis* CdaA_CD_ and GlmM proteins, both of which form dimers in the structures, and of the CdaA_CD_:GlmM_F369_ complex. In the complex structure, a CdaA_CD_ dimer is bound to a GlmM_F369_ dimer in such a manner that GlmM blocks the oligomerization of CdaA_CD_ and formation of active head-to-head cyclase oligomers, thus providing molecular details on how GlmM acts as cyclase inhibitor. The function of a key amino acid residue in CdaA_CD_ in complex formation was confirmed by mutagenesis analysis. As the amino acids at the CdaA_CD_:GlmM interphase are conserved, we propose that the observed inhibition mechanism of CdaA by GlmM is conserved among Firmicutes.

## Introduction

Nucleotide signalling molecules play important roles in helping bacteria to rapidly adapt to changing environmental conditions (1,2). One such signalling nucleotide, cyclic-di-adenosine monophosphate (c-di-AMP), which was discovered a little more than a decade ago (3), plays an important function in the osmotic regulation of bacteria by controlling potassium and osmolyte uptake (4–8). c-di-AMP also plays an important function in regulating cell size, either directly or indirectly through its function in osmotic regulation, cell-wall integrity and susceptibility to beta-lactam antibiotics, which target the synthesis of the peptidoglycan cell wall (9–12).

The function of c-di-AMP has been most extensively studied in a range of Firmicutes bacteria including the Gram-positive model organism *Bacillus subtilis* and Gram-positive bacterial pathogens such as *Staphylococcus aureus, Listeria monocytogenes* and several *Streptococcus* species (9,10,13–17). From these studies, it has become apparent that the cellular level of c-di-AMP needs to be tightly regulated as both an excess and a lack of c-di-AMP can negatively impact bacterial growth, physiology and virulence (17,18). To achieve the optimal level, a dynamic equilibrium must exist between the synthesis of c-di-AMP via diadenylate cyclases and its degradation into 5′-phosphadenylyl-adenosine (pApA) or two molecules of AMP by phosphodiesterases (18–20). As part of the current study, we investigated how the activity the *B. subtilis* c-di-AMP cyclase CdaA is regulated by GlmM, a phosphoglucomutase enzyme required for the synthesis of an essential peptidoglycan precursor.

c-di-AMP is formed from two molecules of ATP by enzymes containing a diadenylate cyclase (DAC) domain. These enzymes have been extensively characterized structurally as well as biochemically, but how their activity is regulated is an aspect that remains poorly understood. *B. subtilis* codes for three diadenylate cyclase enzymes (3,21–24). CdaA (also referred to as DacA in some bacteria) is a membrane-bound cyclase with three predicted N-terminal transmembrane helices and a cytoplasmic catalytic DAC domain. CdaA (DacA) is the “housekeeping” c-di-AMP cyclase in Firmicutes, as it is conserved and often the sole c-di-AMP cyclase in phylum (25,26). The two other *B. subtilis* c-di-AMP cyclases, DisA and CdaS, are soluble proteins, not as widely distributed among bacteria and have more specialized functions, with DisA involved in controlling DNA-repair processes during sporulation or spore outgrowth and CdaS also specifically expressed during the sporulation process (22,23,27). While there are no publications on the 3D-structures of the *B. subtilis* c-di-AMP cyclases, structures are available for the cytoplasmic enzymatic domains of the *L. monocytogenes* and *S. aureus* CdaA/DacA homologs (21,28), the DisA homolog from *Thermotoga maritima* (3) and the CdaS homolog from *B. cereus* (PDB 2FB5). These studies revealed that DAC domains have a mixed αß-fold, with highly conserved DGA and RHR amino acid motifs required for ligand binding (3,21,29). Formation of c-di-AMP requires a head-to-head conformation of two DAC domains. This was first demonstrated in the crystal structure of DisA, a protein which forms an octamer with four DAC domain dimers in the active head-to-head conformation (3). While also the *L. monocytogenes* and *S. aureus* CdaA/DacA catalytic domains and the CdaS protein, where present as dimers and hexamers, respectively, they were in an inactive conformation. These proteins therefore either need to rearrange or more likely form higher oligomers in order to yield active enzymes with DAC domains in the head-to-head dimer conformation (24,28). Recently, another structure of the cytoplasmic catalytic domain of the *L. monocytogenes* CdaA enzyme (Δ100CdaA) has been reported with a c-di-AMP bound between two monomers, which based on the crystal cell packing, were arranged in an active dimer of dimer configuration (30). These findings are consistent with the idea that CdaA (DacA) enzymes will need to form higher oligomers to achieve an active enzyme configuration. Hence, factors influencing the ability of c-di-AMP cyclases to rearrange into an active conformation or to form higher oligomers will be able to regulate the activity of these enzymes.

The genetic arrangement and operon structure coding for the “housekeeping” c-di-AMP cyclase CdaA (DacA) is conserved in Firmicutes (29,31). Two genes, coding for the membrane-linked CdaA regulator CdaR (also named YbbR in some bacteria) and cytoplasmically-located peptidoglycan precursor synthesis enzyme GlmM, are found downstream and in an operon with *cdaA* (29,31). Through recent studies in *B. subtilis, L. monocytogenes, Lactococcus lactis* and *S. aureus,* it is has become apparent that these three genes are not only co-transcribed but that the encoded proteins also from a complex and that CdaR and GlmM can regulate the activity of the c-di-AMP cyclase CdaA (29,32,33). While CdaR has been reported to function as both an activator and repressor of CdaA activity depending on the growth conditions, GlmM has been shown to be a potent inhibitor of the cyclase activity (17,29,31,32,34). However, the molecular mechanisms on how the CdaA cyclase activity is regulated by these proteins are not yet known and this was further investigated as part of this study.

GlmM is a phosphoglucomutase enzyme catalysing the conversion of glucosamine-6-phosphate to glucosamine-1-phosphate, which is subsequently used to produce the essential peptidoglycan precursor UDP-N-acetyl-glucosamine (35). In *B. subtilis, L. monocytogenes* and *L. lactis* a protein-protein interaction between CdaA and GlmM has been detected using bacterial two-hybrid assays performed in *Escherichia coli* (29,31,33). In *B. subtilis* this interaction has been further confirmed by *in vivo* protein cross-linking and pulldown assays (29) and in *L. monocytogenes* by the co-elution of purified proteins (33). The first evidence that GlmM serves as negative regulator of CdaA/DacA activity came from an *L. lactis* strain that produces a GlmM variant that is thought to form a stronger interaction with CdaA; this strain produces lower cellular c-di-AMP levels than the bacteria expressing wildtype GlmM (31). Furthermore, the activity of the soluble recombinant *S. aureus* DacA catalytic domain (DacA_CD_) could be blocked almost completely by the addition of purified GlmM protein in *in vitro* assays and the recombinant proteins were shown to form a stable complex that could be purified via size-exclusion chromatography (28). On the other hand, the activity of GlmM was not affected by the interaction with DacA_CD_ (28). Additional mass-spectrometry and small-angle X-ray scattering data (SAXS) analyses suggested that the complex is composed of a DacA_CD_ dimer and a GlmM dimer (28). Crystal structures of the individual *S. aureus* DacA_CD_ and GlmM dimers revealed that the *S. aureus* DacA_CD_ protein assumed an “inactive” dimer conformation. GlmM had the typical four-domain fold of phosphoglucomutases with a flexible C-terminal domain 4 and the dimer was “M-shaped”, characteristic for this class of enzymes (28). However, a high-resolution structure of the complex could not be obtained and only a model for the complex could be proposed by fitting the individual DacA_CD_ and GlmM dimer structures into the SAXS envelope (28). Based on this, a model was proposed in which GlmM could potentially block the activity of the DacA_CD_ cyclase by preventing the formation of higher oligomers.

Here, we set out to provide atomic resolution information on the CdaA:GlmM complex to gain insight into the molecular mechanism how GlmM can control the activity of the c-di-AMP cyclase enzyme. Using the purified *B. subtilis* CdaA catalytic domain (CdaA_CD_) and purified full-length GlmM or the truncated GlmM_F369_ variant lacking the flexible C-terminal domain 4, we show that the two proteins form a stable complex *in vitro* and that GlmM and GlmM_F369_ are potent inhibits of the cyclase. Crystal structures of the *B. subtilis* CdaA_CD_ cyclase, the GlmM phosphoglucomutase and the CdaA_CD_:GlmM_F369_ complex were obtained, revealing dimer conformations of the individual proteins as well as a dimer of dimer conformation in the complex structure. More importantly, from the complex structure the mechanism by which binding of GlmM inhibits the cyclase activity becomes apparent, that is by preventing the oligomerisation of CdaA and formation of active head-to-head cyclase oligomers.

## Results

### The *B. subtilis* phosphoglucosamine GlmM interacts with and inhibits the activity of the c-di-AMP cyclase CdaA_CD_

Using the purified *S. aureus* DacA_CD_ catalytic domain and GlmM, it has been shown that the proteins form a stable complex *in vitro* and that GlmM is a potent inhibitor of the c-di-AMP cyclase without requiring any additional factors (28). To examine if this is also the case for the *B. sutbilis* proteins, the full-length *B. subtilis* GlmM protein as well as the truncated GlmM_F369_ variant were expressed and purified along with the soluble catalytic domain of the *B. subtilis* c-di-AMP cyclase CdaA_CD_. The GlmM_F369_ variant lacks the flexible C-terminal domain 4 and was constructed to aid subsequent structural investigations. The proteins were expressed as His-tagged proteins in *E. coli* and purified individually via Ni-NTA affinity chromatography followed by size exclusion chromatography (Fig. 1). To test for a CdaA-GlmM interaction, lysates of strains producing CdaA_CD_ and GlmM (Fig. 1A) or CdaA_CD_ and GlmM_F369_ (Fig. 1B) were mixed prior to affinity and size-exclusion chromatography. The elution profiles and analysis of the retention volumes revealed that CdaA_CD_ formed a complex with GlmM and with GlmM_F369_ that eluted as a single, higher-mobility species compared to the individual proteins (Fig. 1). The peak fractions of each complex were further analysed by SDS-PAGE, confirming the presence of both proteins (Fig. 1, inserts). We also determined the binding affinity between GlmM and CdaA_CD_ by microscale thermophoresis (MST). For the MST experiments, increasing concentrations of unlabelled purified GlmM ranging from a final concentration of 0.78 μM to 800 μM were mixed with fluorescence labelled CdaA_CD_ held at a constant final concentration of 25 nM (see experimental procedure sections for details). Based on the thermophoresis and normalized fluorescence change of CdaA_CD_ depending on the GlmM protein concentration a K_d_ of 14.4 μM ± 0.962 was determined (Fig. 1C) indicating a moderate binding affinity. Next, to determine if the *B. subtilis* GlmM protein impacts the activity of CdaA_CD_, *in vitro* cyclase activity assays were performed, and the conversion of ATP (spiked with a small amount of α-^32^P -labelled ATP) into c-di-AMP assessed. The purified *B. subtilis* CdaA_CD_ protein was enzymatically active in the presence of the divalent metal ion Mn^2+^ but showed only limited activity in the presence of Co^2+^ or Mg^2+^ (Fig. 2A) and after 4 h incubation approximately 50% of the ATP substrate was converted to c-di-AMP (Fig. 2B). Addition of GlmM or GlmM_F369_ at a 2:1 molar ratio over CdaA_CD_, led to a significant reduction in the conversion of ATP to c-di-AMP (Fig. 2C). Taken together, these data show that the purified *B. subtilis* CdaA_CD_:GlmM as well as CdaA_CD_:GlmM_F369_ proteins form a stable complex *in vitro* and that both the full-length and truncated GlmM variant, inhibit the activity of the c-di-AMP cyclase CdaA_CD_.

**Figure 1:**
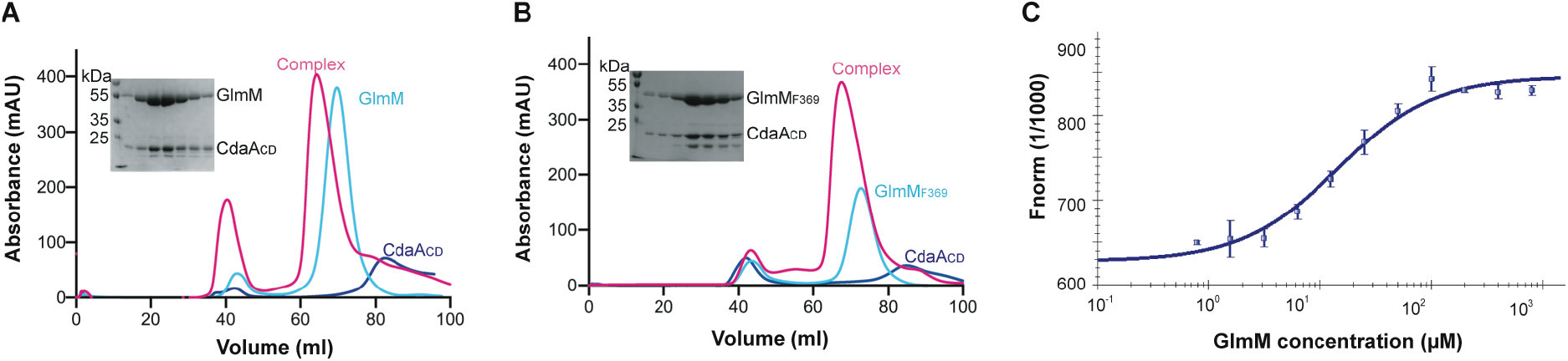
The *B. subtilis* GlmM and CdaA_CD_ proteins form a complex *in vitro*. *A,* FPLC chromatograms and SDS-PAGE gel analysis of the *B. subtilis* CdaA_CD_ and GlmM complex. The *B. subtilis* proteins CdaA_CD_, GlmM and the CdaA_CD_/GlmM complex were purified by Ni-affinity chromatography and size-exclusion chromatography. The FPLC elution profiles, recorded at a wavelength of 280 nm, are shown for CdaA_CD_ (blue), GlmM (cyan) and the CdaA_CD_/GlmM complex (pink). Proteins from the peak fractions of the CdaA_CD_-GlmM complex were separated on a 12% SDS PAGE gel and proteins visualized by Coomassie staining (shown in the insert). *B,* FPLC chromatograms and SDS-PAGE gel analysis of the *B. subtilis* CdaA_CD_ and GlmM_F369_ complex. Same as in (A) but using the C-terminally truncated GlmM_F369_ variant in place of the full-length GlmM protein. The experiments were performed in triplicates and representative chromatograms and gel images are shown. *C*. MST experiment to determine the binding affinity between CdaA_CD_ and GlmM. Increasing concentrations of GlmM were mixed with fluorescently labelled CdaA_CD_ resulted in a gradual change in thermophoresis and fluorescence readings. The normalized fluorescence values multiplied by a factor of 1000 Fnorm (1/1000) were plotted using the NT Analysis Software (NanoTemper Technologies GmbH) to yield the binding curve. The data points are the average values and standard deviations from five independent MST runs. The K_d_ was determined from these data using the NT Analysis Software (NanoTemper Technologies GmbH).

**Figure 2:**
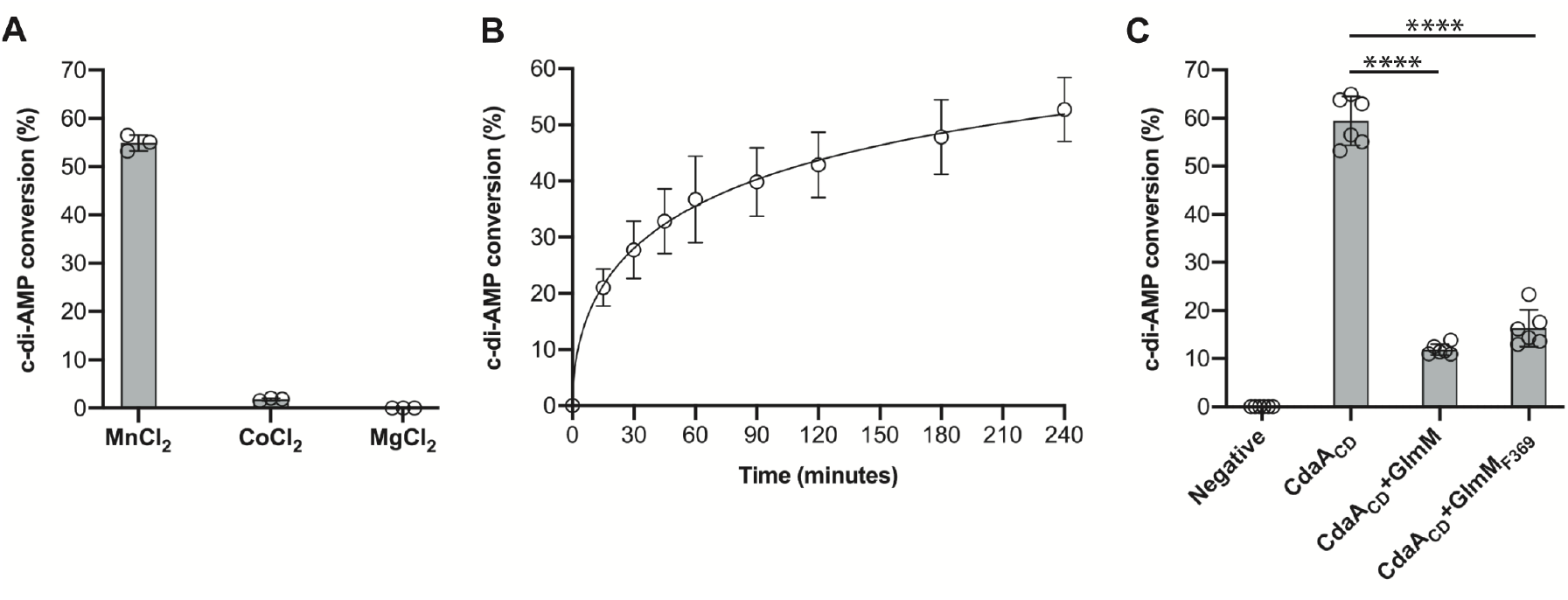
Enzymatic activity of the *B. subtilis* CdaA_CD_ enzyme is inhibited by GlmM or GlmM_F369_. *A,* Metal-dependency of the *B. subtilis* CdaA_CD_ enzyme. The metal-dependency of the *B. subtilis* CdaA_CD_ was assessed by performing enzyme reactions using 5 μM of purified CdaA_CD_ in buffer containing 1 mM Mn^+2^, Mg^+2^ or Co^+2^ ions. After 4 h of incubation, the reactions were stopped, separated by TLC and the percentage conversion of radiolabelled ATP to c-di-AMP determined. The average values and standard deviations (SDs) from three experiments were plotted. *B,* CdaA_CD_ activity time-course experiment. Enzyme reactions were set up with the *B. subtilis* CdaA_CD_ enzyme in buffer containing 1 mM Mn^+2^, aliquots removed, and reactions stopped at the indicated time points and separated by TLC. The percentage conversion of radiolabelled ATP to c-di-AMP was determined and the average values and SDs from three independent experiments plotted. *C,* Enzyme activity of the *B. subtilis* CdaA_CD_.enzyme in the presence of *B. subtilis* GlmM. The enzyme activity of CdaA_CD_ was measured in the absence or presence of GlmM or GlmM_F369_ at a molar ratio of 2:1 GlmM or GlmM_F369_ to CdaA_CD_. After 4 h of incubation, the reactions were stopped, separated by TLC and the percentage conversion of radiolabelled ATP to c-di-AMP was determined. The average values and SDs from six independent experiments plotted. One-way ANOVA tests followed by Dunnett’s multiple comparison were performed to identify statistically significant differences in cyclase activity in the absence or presence of GlmM or GlmM_F369_. *** indicates p<0.0001.

### Crystal structures of the *B. subtilis* CdaA_CD_ and GlmM proteins

To gain atomic level details of the CdaA_CD_ and GlmM protein complex, we started off by determining the crystal structures of the individual proteins. The tag-less *B. subtilis* CdaA_CD_ protein was crystallised and the structure solved at 2.8 Å (Table 1 and Fig. 3). The protein displayed the expected diadenylate cyclase protein fold, with a central β-sheet made up of 6 antiparallel strands flanked by 5 helices (Fig. 3A). However, it lacked the seventh β strand that was seen in the structures of CdaA homologs of other bacteria (21,28). In *B. subtilis* CdaA_CD_ the residues corresponding to this β-strand are instead in a loop that adapts a very similar confirmation to the β strand observed in other CdaA structures. Superposition of the *B. subtilis* CdaA_CD_ structure with the *L. monocytogenes* Δ100CdaA (PDB 4RV7; sequence identity of the full-length proteins is 65%) (21), and *S. aureus* DacA_CD_ (PDB 6GYW; sequence identity of the full-length proteins is 53%) (28) structures, all lacking the N-terminal transmembrane helices, gave r.m.s.ds of 0.79 and 0.75, respectively, highlighting the overall structural similarities of these enzymes (Fig. 3B). The *B. subtilis* CdaA_CD_ structure was solved as a dimer in the asymmetric unit with hydrogen-bonding interactions observed at the interaction interface (Fig 3C). Interactions were observed between the side chains of amino acid residues Asn166, Thr172 and Leu174 (site 1) and residues Leu150, Lys153 and Met155 (site 2) (Fig 3C). Similar hydrogen-bonding interactions were also identified in the *S. aureus* DacA_CD_ and *L. monocytogenes* Δ100CdaA structures with amino acid residues in site 1 being absolutely conserved (28,30) (Fig. S1). Analysis of the interface with PDBePISA (36) indicated a buried surface of 1400 Å^2^, which is similar to the value of 1460 Å^2^ previously reported for the *S. aureus* DacA_CD_ protein, indicative of a stable dimer formation. In this dimer confirmation, the active sites face opposite directions and hence cannot be engaged in a catalytically active head-to-head conformation (Fig. 3A). Taken together, these data indicate that the conformationally-inactive dimerization interface is conserved among different CdaA homologs in Gram-positive bacteria and that the enzyme needs to form higher oligomers for catalysis.

**Table 1:**
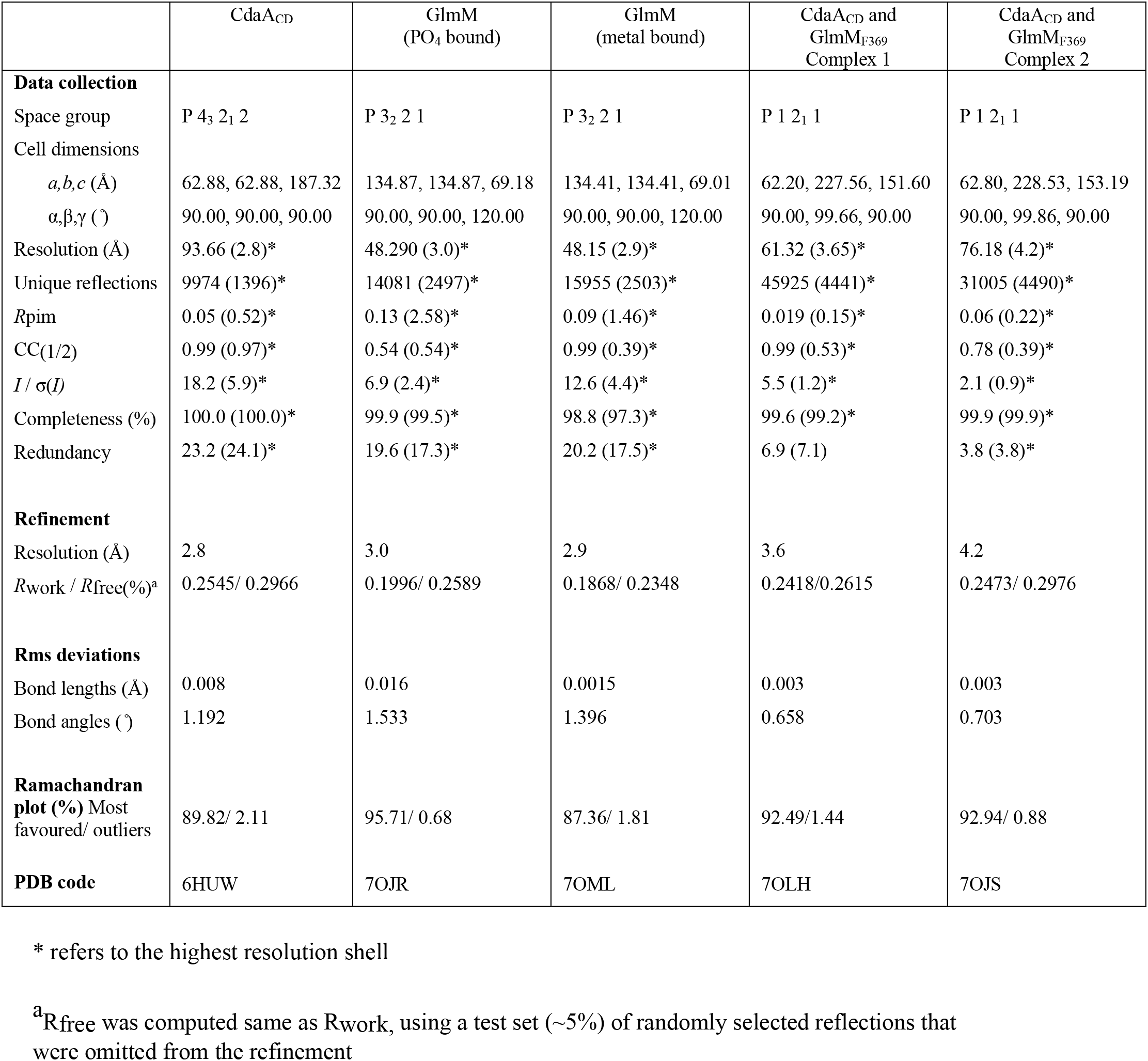
Crystallographic data and refinement statistics

**Figure 3:**
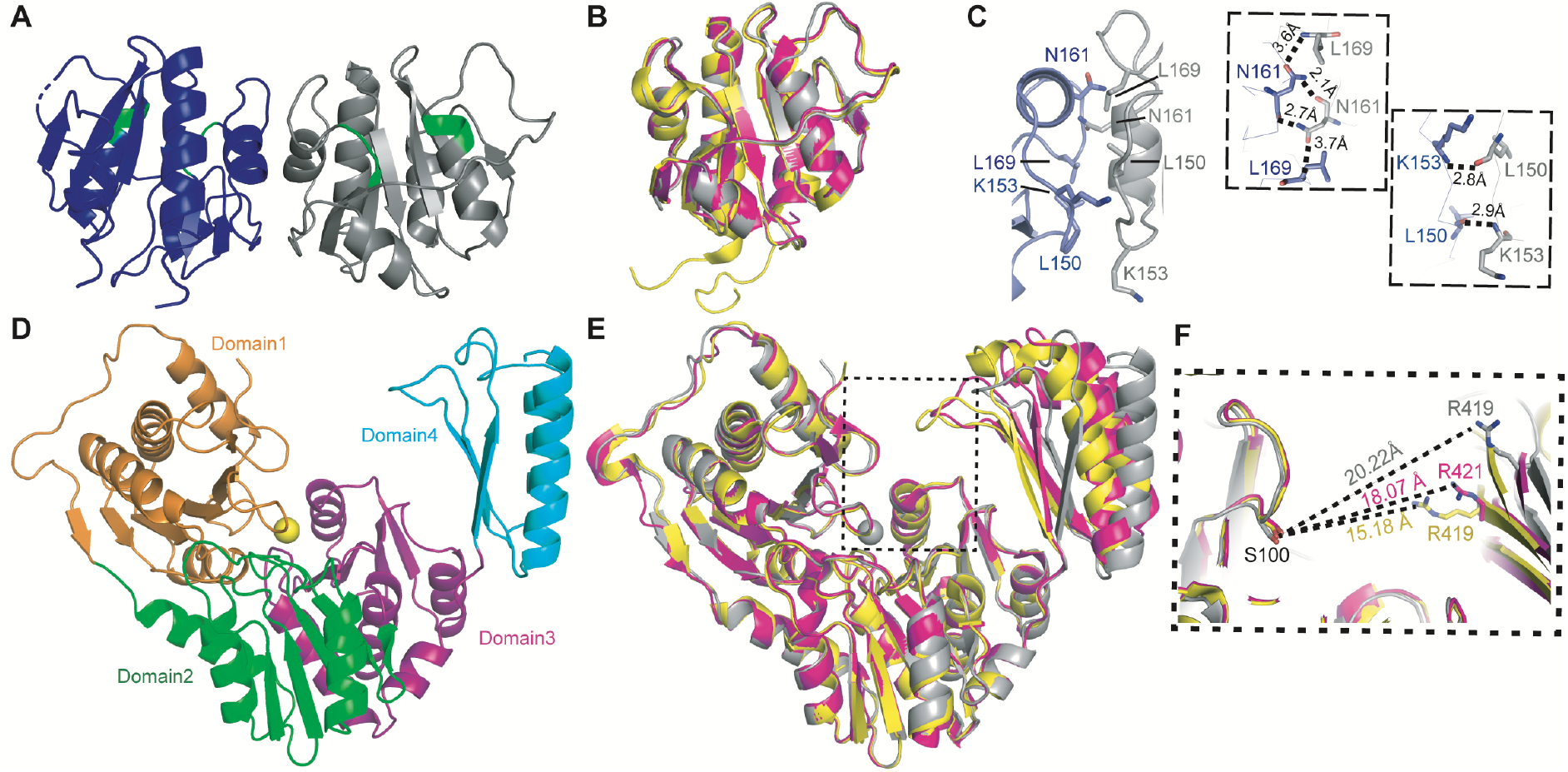
Crystal structures of the *B. subtilis* CdaA_CD_ and GlmM enzymes. *A, B. subtilis* CdaA_CD_ structure in cartoon representation. The CdaA_CD_ protein crystallized as dimer and individual CdaA_CD_ monomers are shown in blue and grey, with active site DGA and RHR motifs highlighted in green. *B,* Superimposition of the *B. subtilis, L. monocytogenes* and *S. aureus* CdaA_CD_ (DacA_CD_) structures. Monomers of the *B. subtilis* CdaA_CD_ (grey), *L. monocytogenes* CdaA_CD_ (yellow; PDB ID 4RV7) and *S. aureus* DacA_CD_ (pink; PDB ID 6GYW) protein structures were superimposed using COOT (48). *C,* CdaA_CD_ dimer interface and zoomed in view showing residues Leu150, Lys153, Asn161 and Leu169, which form hydrogen-bonds, in stick representation. *D*, *B. subtilis* GlmM structure in cartoon representation. The GlmM protein crystalized as a dimer (Fig. S2) with one monomer in the asymmetric unit. The GlmM monomer displayed the typical 4 domain architecture, and the individual domains are represented in different colours. A metal-ion was bound to the protein and is shown as yellow sphere. *E,* Superposition of the *B. subtilis, S. aureus* and *B. anthracis* GlmM structures. *B. subtilis* GlmM (grey), *B. anthracis* GlmM (yellow; PDB 3PDK) and *S. aureus* GlmM (pink; PDB 6GYX) monomer structures were superimposed using COOT (48). *F,* Zoomed in view of the superimposed *B. subtilis* (grey; metal-ion bound structure)*, S. aureus* (pink) and *B. anthracis* (yellow) GlmM structures showing the distances between the indicated Arg residue within the phosphate binding site and the active site Ser residue. Images were generated using PyMOL (The PyMOL Molecular Graphics System, Version 2.0 Schrödinger, LLC).

The His-tagged *B. subtilis* GlmM protein was crystallised and the structure solved by molecular replacement using the *B. anthracis* GlmM structure (PDB 3PDK; (37)) as the search model (Table 1 and Fig. 3D). The *B. subtilis* GlmM protein displayed a four-domain architecture typical for phosphoglucosamine mutase proteins (28,37) (Fig. 3D). Domains 1-3 are comprised of α-β mixed cores linked via a flexible loop to domain 4, which displays a 3-stranded β sheet fold surrounded by two α-helices (Fig. 3D). While one GlmM molecule was present in the asymmetric unit, the typical “M-shaped” GlmM dimer arrangement was observed in the crystal cell packing (Fig. S2). Interactions were formed between domains 1, leading to the formation of a large groove at the top of the dimer molecule, mostly formed by domain 2 and the active site of each monomer subunit facing the opposite direction. Two different structures were solved for the *B. subtilis* GlmM protein at 2.9 and 3.0 Å resolutions with a superposition r.m.s.d. score of 0.29 (Table 1 and Fig. S2). One of the crystal structures was obtained with a divalent cation bound to the catalytic serine residue, which during catalysis is thought to be converted to a phosphoserine residue and the metal ion playing an important role during catalysis (Fig. S2). The exact type of metal ion could not be deduced due to the limitation of the structural resolution. However, we speculate that it is a magnesium ion, as magnesium was present in the crystallisation conditions and this metal ion is usually also bound in fully active enzymes. Furthermore, when a magnesium ion was modelled into the structure and analyzed using the program CheckMyMetal (38), a better fit was observed as compared to zinc or calcium ions, which could also fill the density. In the second structure, a phosphate molecule (PO4) was bound to Arg419, located within a loop region in domain 4 (Fig S2B) at a similar location as observed in the *B. anthracis* GlmM structure (37). The superimposition of the *B. subtilis* GlmM structure with the *S. aureus* (6GYZ; (28)) and *B. anthracis* (3PDK; (37)) GlmM structures, gave small r.m.s.d. values of 1.0261 and 1.0668, respectively (Fig. 3E), indicating high similarity. However, the inter-residue distance between Arg419 in the phosphate binding site in domain 4 and the catalytic Ser100 in domain 1 was 20.22 Å in the phosphate bound *B. subtilis* GlmM structure compared to 18.4 Å in the *S. aureus* GlmM (PDB 6GYZ) or 15.18 Å in the *B. anthracis* GlmM (PDB 3PDK) structures (Fig. 3F). This highlights the flexibility of domain 4 in GlmM enzymes and also reveals that the *B. subtilis* GlmM protein was captured in most open state of the enzyme reported so far in a crystal structure.

### Structure of the *B. subtilis* CdaA_CD_:GlmM_F369_ complex

To understand how GlmM interacts and inhibits the CdaA, we next aimed to obtain the structure of the complex. Any crystals obtained for the *B. subtilis* CdaA_CD_:GlmM complex diffracted poorly. On the other hand, diffracting crystals were obtained for the CdaA_CD_:GlmM_F369_ complex, in which the GlmM protein lacks the flexible C-terminal domain 4. The crystals were obtained under two different conditions (see experimental procedures section), and the structure of the CdaA_CD_:GlmM_F369_ complex could be solved at 3.6 Å (Complex 1) and at 4.2 Å (Complex 2) by molecular replacement using the *B. subtilis* CdaA_CD_ and GlmM (dimer) structures as search models (Table 1 and Fig. 4, Fig. S3). While obtained under two different conditions, complex 1 and complex 2 were nearly identical and overlapped with an r.m.s.d of 0.22 Å (Fig. S4A). Furthermore, in both complex structures, three complex molecules were obtained in the asymmetric unit and each complex was composed of a GlmM_F369_ dimer and a CdaA_CD_ dimer in the inactive dimer configuration (Fig. 4 and Fig. S3). The three complexes obtained in the asymmetric unit were almost identical to each other, as indicated by the superposition r.m.s.d. of 0.15 Å to 0.20 Å for complex 1 (Fig. S4B) and of 0.15 Å to 0.16 Å for complex 2 (Fig. S4C). Since the obtained complex structures were basically identical, all further descriptions are based on the higher resolution complex 1 structure. In the complex, a CdaA_CD_ dimer was positioned in the large groove at the top of the GlmM_F369_ dimer and formed interactions with domain 2 of GlmM (Fig 4A-4C). The complex was asymmetric, with one of the CdaA_CD_ monomer, CdaA_CD_(2) (shown in light blue in Fig. 4) placed in the center of the GlmM_F369_ groove and the other monomer, CdaA_CD_(1) (shown in dark blue in Fig. 4) projecting towards the solvent. Similarly, most of the interactions of the GlmM_F369_ dimer with the CdaA_CD_ dimer were made by GlmM_F369_(1) (shown in dark pink in Fig 4). PDBePISA analysis revealed an average buried surface area of 996 Å^2^ in the interface between GlmM_F369_(1) and the CdaA_CD_ dimer, which was stabilized by 4 hydrogen bond and 4 ionic bond interactions between GlmM_F369_(1) and CdaA_CD_(1) and 5 hydrogen bond and 3 ionic bond interactions with CdaA_CD_(2) (Table S1 and Fig. S5). On the other hand, only an average 220.3 Å^2^ surface area is occluded in GlmM_F369_(2) (shown in light pink in Fig. 4). Based on the PDBePISA analysis, GlmM_F369_(2) only formed two hydrogen bond interactions with the CdaA_CD_(2) monomer but no interaction with CdaA_CD_(1) (Table S1 and Fig. S5). A more detailed analysis of the interface showed that several interactions are made between two α-helices from domain 2 of GlmM_F369_(1), α1 and α2, with the CdaA_CD_(1) and CdaA_CD_(2) monomers, respectively (Fig 4C and 4D). The main interactions in the complex were formed between three residues, D151, E154, and D194 of domain 2 in GlmM_F369_(1) and residue R126 in each of the CdaA_CD_ monomers. More specifically, ionic bonds were formed between residue D195 in GlmM_F369_(1) and residue R126 in CdaA_CD_(2). In addition, salt bridges were formed between residues D151 and E154 in GlmM_F369_(1) and residue R126 but this time from CdaA_CD_(1) (Fig 4C and 4D, Table S2 and Fig. S5). The data suggest that residue R126 in CdaA_CD_ is potentially one of the most critical residues for complex formation, as it contributes to a number of ionic as well as hydrogen-bond interactions and even though the complex is asymmetric, it contributes to interactions in both CdaA_CD_ monomers.

**Figure 4:**
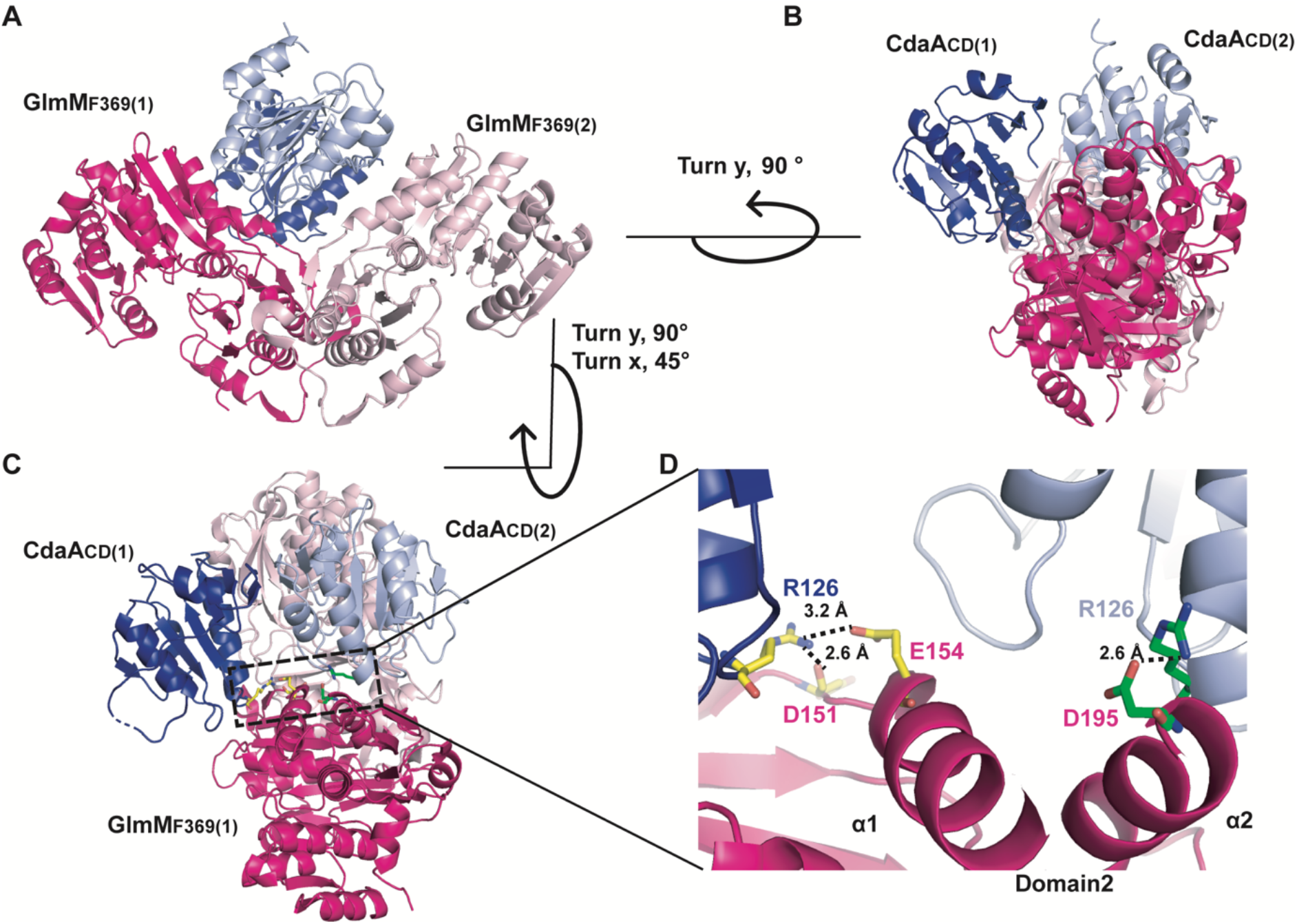
Crystal structure of the *B. subtilis* CdaA_CD_:GlmM_F369_ complex 1. (*A-D*) Structure of the *B. subtilis* CdaA_CD_:GlmM_F369_ complex 1 shown in cartoon representation. The *B. subtilis* CdaA_CD_:GlmM_F369_ complex crystallized as a dimer of dimers with individual GlmM_F369_ monomers shown in dark pink [GlmM_F369_(1)] and light pink [GlmM_F369_(2)] and individual CdaA_CD_ monomers shown in dark blue [CdaA_CD_(1)] and light blue [CdaA_CD_(2)], respectively. The complex is show in *A,* in front view, *B,* in side view (rotated 90º along the y-axis) and *C*, in top-side view rotated at the angle as indicated with respect to (A). *D,* a zoom in view of the CdaA_CD_/GlmM_F369_ interface is shown. Residue Arg126 from CdaA_CD_(1) forms H-bond and ionic interactions with Asp 151 and Glu154 of GlmM_F369_(1) (residues shown in yellow), and residue Arg126 from CdaA_CD_(2) forms ionic interactions with Asp195 in GlmM_F369_(1) (residues shown in green). The images were prepared in PyMOL (The PyMOL Molecular Graphics System, Version 2.0 Schrödinger, LLC).

### CdaA_CD_ cannot form active oligomers in complex with GlmM

To gain insight how GlmM inhibits the activity of the c-di-AMP cyclase, we inspected the location of the active sites of CdaA_CD_ in the complex. The active site of DAC-domain enzymes is characterized by DGA and RHR motifs, corresponding to residues D_171_GA and R_203_HR in *B. subtilis* CdaA (Fig. 5; areas highlighted in yellow and green in the CdaA_CD_ monomers). The active site in CdaA_CD_(2) was completely occluded upon interaction with the GlmM_F369_ dimer (Fig. 5, dark blue CdaA_CD_ monomer with active site region highlighted in yellow) but the active site in CdaA_CD_(1) appeared at least partially exposed (Fig. 5; light blue CdaA_CD_ monomer with active site region highlighted in green). For CdaA_CD_ to produce c-di-AMP active head-to-head dimers need to be formed (3). The crystal structure of the *L. monocytogenes* Δ100CdaA cyclase was recently determined with a c-di-AMP molecule bound in the catalytic site and an active head-to-head dimer conformation seen in the crystal packing (30). Using the *L. monocytogenes* Δ100CdaA structure (PDB 6HVL) as model, an active *B. subtilis* CdaA_CD_ dimer was modelled and superimposed on CdaA_CD_(1) in the complex structure (Fig. 5). Although the active site of the CdaA_CD_(1) was exposed and accessible in the complex with GlmM_F369_, in an active dimer conformation, parts of the second CdaA_CD_ molecule would collide and overlap with GlmM_F369_, highlighting that also CdaA_CD_(1) cannot form active head-to-head oligomers in the complex (Fig. 5). Taken together, these data indicate that in the complex, the interaction of GlmM with CdaA_CD_ will prevent the formation of functional diadenylate cyclase enzyme oligomers, which is essential for the formation of c-di-AMP. The crystal structure of the CdaA_CD_:GlmM_F369_ complex therefore provides insight on an atomic level on the catalytic inhibition of the diadenylate cyclase CdaA_CD_ by the phosphoglucosamine enzyme GlmM.

**Figure 5:**
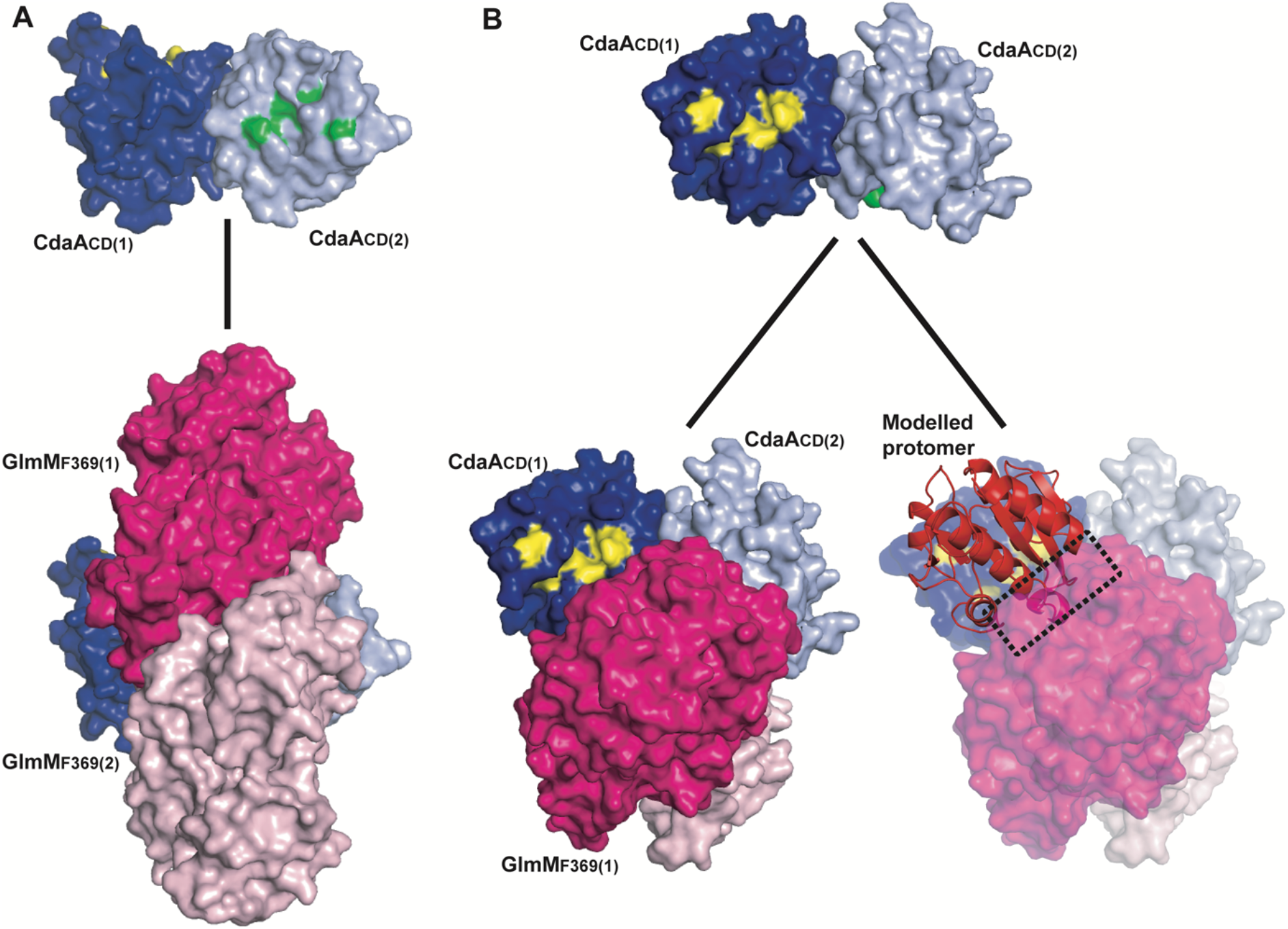
Formation of an active CdaA_CD_ dimer is blocked in the complex. (*A-B*) Structure of the *B. subtilis* CdaA_CD_ dimer from complex 1 (top panels) and the complete CdaA_CD_:GlmM_F369_ complex 1 (bottom panels) shown as space filling models. The active site DGA and RHR motifs in CdaA_CD_(1) (dark blue protomer) are shown in yellow and in CdaA_CD_(2) (light blue protomer) in green. *A*, Bottom panel, complex model showing that the active site (green residues) in CdaA_CD_(2) (light blue protomer) are completely occluded in the complex and *B,* the active site (yellow residues) of CdaA_CD_(1) (dark blue protomer) remains partially exposed. However, the modelling of another CdaA_CD_ protomer (in red) that forms an active dimer with CdaA_CD_(1) (dark blue protomer) results in protein-protein clashes with the GlmM_F369_(1) monomer in the complex. The active CdaA_CD_ dimer was modelled based on the *L. monocytogenes* Δ100CdaA (PDB 6HVL) structure (30).

### Small angle X-ray scattering analysis of *B. subtilis* CdaA_CD_:GlmM complex

To determine whether the full-length *B. subtilis* GlmM protein interacts with CdaA_CD_ in a similar manner as observed for GlmM_F369_, a structural characterisation of the CdaA_CD_:GlmM complex was performed via small-angle X-ray scattering (SAXS). To this end, the individual purified *B. subtilis* CdaA_CD_ and GlmM proteins as well as the purified CdaA_CD_:GlmM complex and as control the CdaA_CD_:GlmM_F369_ complex, were passed over an analytical size-exclusion column, followed by continuous automated SAXS data collection throughout the run (Fig. 6, Fig S6 and Table S2). For CdaA_CD_ and GlmM, the reconstructed maps were consistent with the proteins forming dimers and the maps were a good fit for the *B. subtilis* CdaA_CD_ dimer (Fig. 6A) and GlmM dimer (Fig. 6B) structures, respectively. The reconstructed map for the CdaA_CD_:GlmM complex (V_c_: 890.1, R_g_: 44.65 Å and d_max_: 161 Å) was bigger in volume and dimensions as compared to the individual maps calculated for GlmM V_c_: 625.8; R_g_: 37.29 Å, d_max_: 122 Å) and CdaA_CD_ (V_c_: 373.2, R_g_: 26.94 Å, d_max_: 88 Å), which is consistent with the formation of a complex. From the Guinier plot analysis, the molecular weight of the *B. subtilis* CdaA_CD_:GlmM complex was calculated to be 130 kDa, which is consistent with the theoretical molecular weight of 144.46 kDa for a complex made of two CdaA_CD_ and two GlmM molecules. To fit a CdaA_CD_:GlmM dimer complex into the reconstructed map, a model of the complex with full-length GlmM was first constructed by superimposing the crystal structure of full-length GlmM onto the CdaA_CD_:GlmM_F369_ complex structure. The resulting complex model was subsequently fitted in the reconstituted SAXS envelope of the complex. A good fit of the CdaA_CD_:GlmM dimer model complex into the reconstructed envelope was observed, however an elongated density on one side remained unoccupied (Fig. 6C). It is plausible that the flexible C-terminal domain 4 of the GlmM protein is responsible for this extra density. As control, a SAXS experiment was also performed using the CdaA_CD_:GlmM_F369_ complex sample for which the X-ray structure was obtained. The dimensions of the CdaA_CD_:GlmM_F369_ complex were V_c_: 656.8, R_g:_ 37.51 Å and d_max_: 117.5 Å and the molecular weight was calculated to be 97.5 kDa, which is consistent with the theoretical molecular weight of 120 kDa for a complex made of two CdaA_CD_ and two GlmM_F369_ molecules. Similarly, a good fit of the CdaA_CD_:GlmM_F369_ dimer complex structure was obtained when fitted into the reconstructed SAXS envelope data (Fig 6D). These data suggest that the full-length GlmM protein likely forms a dimer-of-dimer complex with the c-di-AMP cyclase CdaA_CD_ and might assume a similar arrangement as observed for the CdaA_CD_:GlmM_F369_ complex.

**Figure 6:**
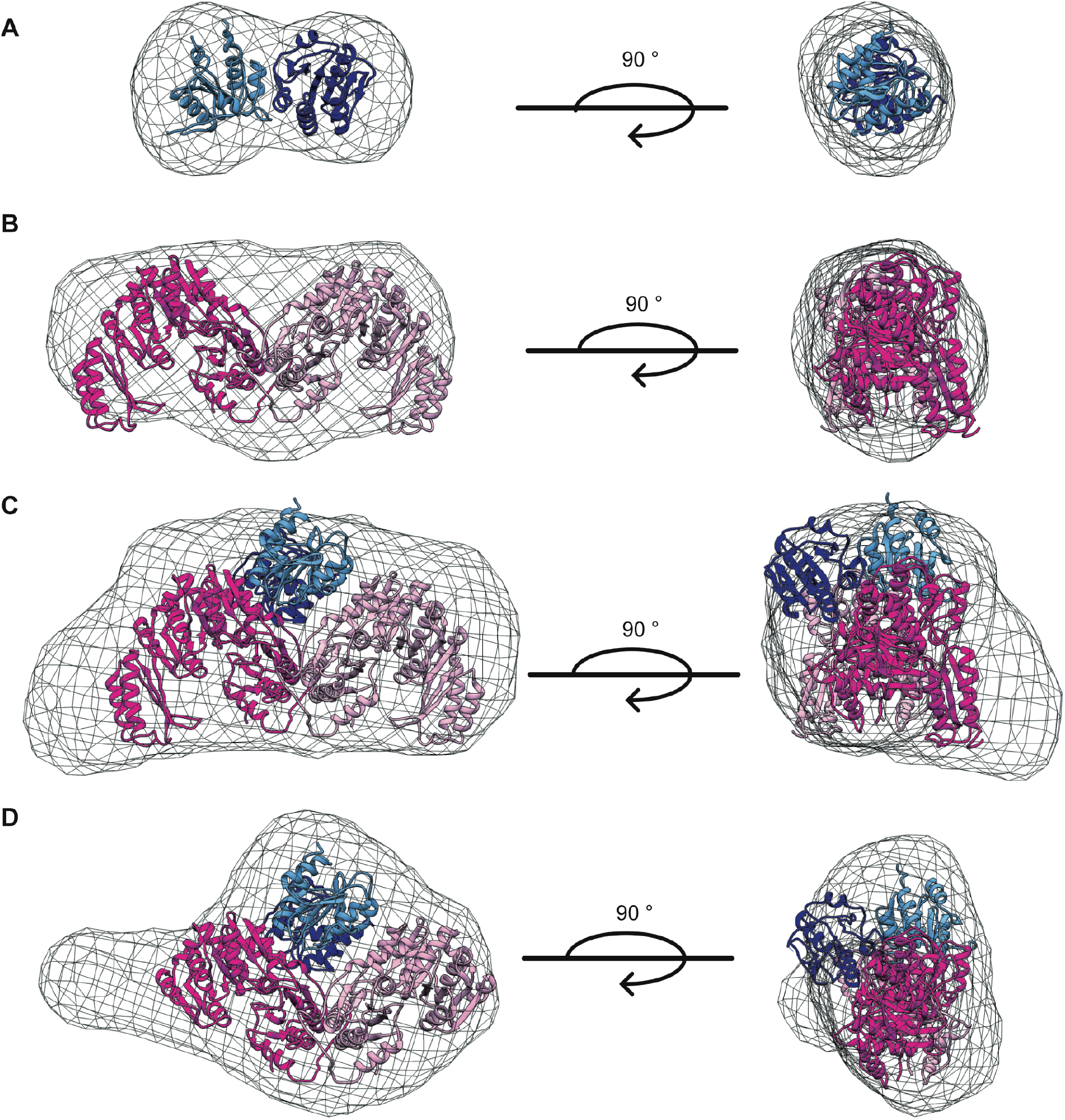
Small Angle X-ray Scattering (SAXS) data of the *B. subtilis* CdaA_CD_, GlmM proteins and the CdaA_CD_:GlmM and CdaA_CD_:GlmM_F369_ complexes. *A-D,* SAXS envelopes with fitted protein structures. For the SAXS experiment, 50 μl of CdaA_CD_ (5 mg/ml), GlmM (24 mg/ml) and the complex CdaA_CD_:GlmM (24 mg/ml) and the CdaA_CD_:GlmM_F369_ complex (10 mg/ml) were injected onto a high pressure Shodex column coupled to the B21 Small-Angle X-Ray beamline at the Diamond Light Source (Didcot, UK). The data analysis and envelope reconstruction were performed using ScÅtter (51). The program Chimera (53) was used to visualise the reconstructed envelopes as maps into which the protein structures were fitted. *A,* SAXS envelope of the *B. subtilis* CdaA_CD_ protein with a CdaA_CD_ dimer protein structure fitted into the envelope. *B,* SAXS envelope of the *B. subtilis* GlmM protein with the GlmM protein dimer structure fitted into the envelope. *C,* SAXS envelope of the *B. subtilis* CdaA_CD_:GlmM complex with a CdaA_CD_:GlmM dimer model structure fitted into the envelope. *D*, SAXS envelope of the *B. subtilis* CdaA_CD_:GlmM_F369_ complex with the structure of CdaA_CD_:GlmM_F369_ dimer complex fitted into the envelope.

### Arginine 126 in *B. subtilis* CdaA_CD_ is essential for complex formation

The complex structure highlighted key interactions between residues D194 and residues D151/E154 in GlmM with residue R126 in each of the CdaA_CD_ monomers (Fig. 5D). To confirm our structural findings, a site-directed mutagenesis analysis was performed. To this end, D195A, D151A/E154A and D151A/E154A/D191A alanine substitution GlmM variants were created. Furthermore, residue R126 in CdaA_CD_, which in both monomers makes contacts with GlmM, was mutated to an alanine. The different alanine substitution variants were expressed and purified from *E. coli* and complex formation assessed by size-exclusion chromatography. While our initial experiments using the GlmM single, double and triple alanine substitution variants appeared not or only marginally to affect complex formation with CdaA_CD_ (Fig. S7), no complex-specific peak was observed when the interaction between the CdaA_CD_-R126A variant and GlmM was assessed. Instead, two peaks were observed for the CdaA_CD_-R126A:GlmM sample, one corresponding to the retention volume of GlmM and the another to CdaA_CD_-R126A (Fig. 7A). Analysis of the elution fractions from the CdaA_CD_-R126A/GlmM sample by SDS-PAGE and Coomassie staining showed that only a very small fraction of the CdaA_CD_-R126A protein co-eluted with GlmM (Fig. 7A). These data highlight that, consistent with the structural data, residue R126 in CdaA_CD_ plays a key role for the complex formation with GlmM. Based on these data, it can be predicted that the cyclase activity of the CdaA_CD_-R126A variant should now longer be inhibited by GlmM. To test this experimentally, *in vitro* cyclase enzyme activity assays were performed. The CdaA_CD_-R126A variant was active, although the activity was reduced as compared to wild-type CdaA_CD_ (Fig. 7B). Importantly and in contrast to wild-type CdaA_CD_, the enzyme activity of this variant was no longer inhibited by the addition of GlmM (Fig. 7B). These data show that residue Arg126 in *B. subtilis* CdaA_CD_ plays a critical role for complex formation and that GlmM can only inhibit the activity of the c-di-AMP cyclase after the formation of a stable complex.

**Figure 7:**
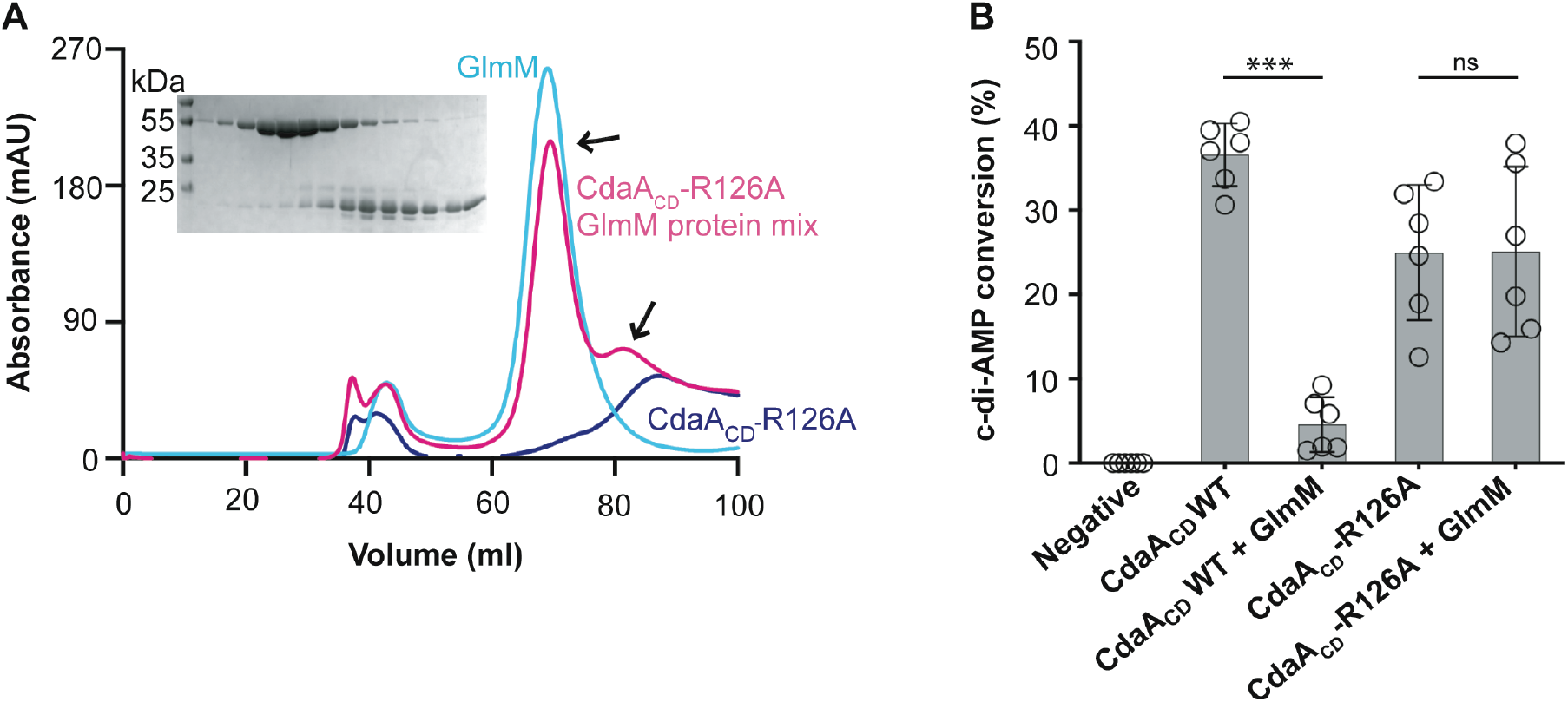
Residue Arg126 in *B. subtilis* CdaA_CD_-R126A is important for complex formation with GlmM. *A,* FPLC chromatograms and SDS-PAGE gel analysis of the *B. subtilis* CdaA_CD_-R126A, GlmM and a mix of both proteins. The *B. subtilis* CdaA_CD_-R126A, GlmM and the CdaA_CD_-R126A/GlmM protein mixture were purified by Ni-affinity chromatography and size-exclusion chromatography. The FPLC elution profiles, recorded at a wavelength of 280 nm, are shown for CdaA_CD_-R126A (blue), GlmM (cyan) and the CdaA_CD_-R126A/GlmM mixture (pink). Proteins from the peak fractions of the CdaA_CD_-R126A/GlmM sample were separated on a 12% SDS PAGE gel and proteins visualized by coomassie staining (shown in the insert). The experiments were performed in triplicates and representative chromatograms and gel images are shown. *B,* Enzymatic activity of wild type CdaA_CD_ and the CdaA_CD_-R126A protein variant in the absence or presence of GlmM. Enzyme activity assays were performed with 5 μM purified CdaA_CD_ or CdaA_CD_-R126 in absence or presence of 10 μM GlmM protein. After 4 h of incubation, the reactions were stopped, separated by TLC and the percentage conversion of radiolabelled ATP to c-di-AMP determined. The average values and SDs from six experiments were plotted. T-tests were performed to determine statistically significant differences in enzyme activity of CdaA_CD_ or CdaA_CD_-R126A in the absence or presence of GlmM. ns indicates not statistically significant; *** indicates p<0.001.

## Discussion

In this study, we show that the *B. subtilis* GlmM and CdaA_CD_ cyclase domain form a stable dimer-of-dimer complex. GlmM acts through this protein-protein interaction as a potent inhibitor of the c-di-AMP cyclase without requiring any additional factors. Based on the atomic-resolution complex structure data, we suggest that GlmM inhibits the activity of CdaA_CD_ by preventing the formation of active head-to-head cyclase oligomers.

For CdaA to produce c-di-AMP, two monomers need to be arranged in an active head-to-head conformation. As part of this study, we determined the structure of the *B. subtilis* CdaA_CD_ protein and show that it has the typical DAC domain fold. While the protein was also found as a dimer in the structure, the dimer was in an inactive conformation, with the two active sites facing in opposite directions. The interface creating the inactive dimer conformation is conserved among CdaA proteins. The *L. monocytogenes* and *S. aureus* homologs, for which structures are available, were found in the same inactive dimer conformation even though the proteins crystallized under different conditions and were found in different space groups (21,28,30). This makes it less likely that a crystallographic symmetry artefact is responsible for the observed inactive dimer configuration. In addition to the inactive dimer configuration within an asymmetric unit, an active dimer conformation was observed in the *L. monocytogenes* Δ100CdaA protein by inspecting adjacent symmetry units (30). However, no such active dimer head-to-head conformations were identified for the *B. subtilis* CdaA_CD_ protein across different symmetry units in the current structure. While not further investigated as part of this study, previous work on the *S. aureus* homolog indicated that the inactive dimer conformation is very stable, and in order for the protein to produce an active enzyme, the protein needs to form higher-level oligomers (28). Given the similarity in the interaction interface, this is likely also the case for the *B. subtilis* CdaA_CD_ enzyme and we would suggest that the *B. subtilis* CdaA_CD_ dimer observed in the structure is unlikely to rearrange into an active dimer conformation.

We also solved the structure of the *B. subtilis* GlmM enzyme. The protein assumed the typical 4-domain architecture previously reported for GlmM enzymes (28,37) and the “M shape” in the dimer conformation, which in the case of the *B. sutbtilis* GlmM protein was formed across two adjacent crystallographic units. The *B. subtilis* GlmM structure further highlighted the flexibility of the most C-terminal domain 4, which was found in the most open conformation seen in any GlmM protein structure up to date. The conformational flexibility of domain 4 is probably also a main factor why we were unsuccessful in determining the structure of a complex between CdaA_CD_ and full-length GlmM. However, GlmM domain 4 is not required for the interaction with and inhibition of CdaA_CD_, since a *B. subtilis* GlmM variant lacking domain 4 formed a complex and inhibited the activity of *B. subtilis* CdaA_CD_. Futhermore, by using the *B. subtilis* GlmMF _F369_ variant lacking the flexible domain 4, we were able to obtain the structure of the CdaA_CD_:GlmM_F369_ complex, revealing for the first time structural details at the atomic level for this complex, thereby identifying the amino acids important for the interaction between the two proteins. Several electrostatic interactions were detected between CdaA_CD_:GlmM_F369_: between the negatively charged residues D151, E154 and D195 in domain 2 of *B. subtilis* GlmM_F369_ with the positively charged residue R126 in CdaA_CD_. We could further show that replacing residue R126 in CdaA_CD_ with an alanine abolished complex formation and the activity of the CdaA_CD_-R126 variant was no longer inhibited by GlmM. A direct protein-protein interaction between CdaA and GlmM has now been reported for these proteins in several Firmicutes bacteria, and hence the amino acids required for the interaction might be conserved. Indeed, a ConSurf (39) analysis using 250 CdaA protein sequences, showed that the residue corresponding to R126 in *B. subtilis* CdaA is conserved between the different homologs (Fig. S8). Likewise, all the three negatively charged residues, D151, E154 and D195 in GlmM, which mediate the primary electrostatic interactions with R126 of CdaA_CD_, are highly conserved (Fig. S8). In previous work, we have shown, that the *S. aureus* DacA_CD_ (the CdaA_CD_ homolog) does not interact with GlmM proteins from *E. coli* and *Pseudomonas aeruginosa,* two Gram-negative bacteria (28). While negatively charged amino acids corresponding to residues E154 and D195 in *B. subtilis* GlmM are also found in the GlmM protein from the Gram-negative bacteria (D153 and E194 in *E. coli* and D152 and E193 in *P. aeruginosa*), the amino acids at the equivalent position of D151 in *B. subtilis* GlmM is an arginine residue (R150 in *E. coli* and R149 in *P. aeruginosa*), which may hinder the complex formation between CdaA (DacA), and GlmM proteins of Gram-negative bacteria (28).

In a previous work, we proposed a model whereby GlmM inhibits the activity of CdaA cyclase activity by preventing the formation of active head-to-head oligomers (28). In the absence of an actual atomic resolution structure of the complex, this model was based on SAXS envelope data and fitting individual protein structures. The model predicted that the likely interaction site between CdaA and GlmM proteins is domain 2 of GlmM (28). The structure of *B. subtilis* CdaA_CD_:GlmM_F369_ complex we present here now provides experimental evidence for such a model and shows that GlmM indeed inhibits the activity of the CdaA cyclase *in vitro* by preventing the formation of active head-to-head oligomers. The K_d_ between CdaA_CD_ and GlmM was in the μM range, which will likely allow complex formation and dissociation in response to changes in protein levels and/or changes in cellular or environmental conditions. GlmM is an essential metabolic enzyme required for the synthesis of the peptidoglycan precursor glucosamine-1-P and thought to be predominantly located within the cytoplasm of the cell (33). However based on the work presented in this study and previous findings, it is assumed that under certain conditions, a fraction of GlmM will localize to the bacterial membrane and interact with and inhibit the activity of the membrane-linked c-di-AMP cyclase CdaA (DacA) (28,32,33). As a result of this interaction, cellular c-di-AMP levels would decrease and consequently potassium and osmolyte uptake increase. Recent work on *L. monocytogenes* suggests that GlmM regulates CdaA during hyperosmotic stress conditions, as during these conditions, overexpression of GlmM has been shown to result in a decrease in cellular c-di-AMP levels (33). The resulting activation of potassium and osmolyte transporter due to a drop in cellular c-di-AMP levels will help cells to counteract the water loss under osmotic stress conditions and aid in bacterial survival. However, what exact cellular changes caused by the osmotic upshift lead to a relocalization of GlmM to the membrane to form a complex with CdaA is currently not known and will require further investigation.

The level of c-di-AMP is regulated by a fine balance between the activities of the cyclase, which synthesizes c-di-AMP, and the phosphodiesterases, which break it down. Interestingly, these two classes of enzyme appear to be regulated very differently; whereas the activity of several phosphodiesterases has been shown to be regulated by small molecules, cyclase activity appears to be regulated through protein-protein interaction. For example, the stringent response alarmone (p)ppGpp has been shown to inhibit the activity both GdpP and PgpH enzymes (11,20). Furthermore, binding of heme to the Per-ARNT-Sim (40) signalling domain in GdpP (which is separate from its DHH/DHHA1 enzymatic domain that is responsible for the degradation of c-di-AMP) was shown in *in vitro* enzyme assays to lead to reduced phosphodiesterase activity (40). Interestingly, the ferrous form of heme bound to GdpP could form a pentacoordinate complex with nitric oxide (NO), resulting in increased c-di-AMP phosphodiesterase activity. Based on these data it has been suggested that GdpP is a heme or NO sensor, resulting in decreased or increased activity respectively (41). The function consequence and impact of (p)ppGpp, heme or NO binding to the phosphodiesterases on bacterial physiology has not yet been fully investigated. However, from these data it is clear that the activity of the c-di-AMP phosphodiesterases can be regulated by small molecule ligands.

On the other hand, several proteins have been found to interact with and regulate the activity of c-di-AMP cyclases. The *B. subtilis* DisA protein is involved in monitoring the genomic stability ensuring that damaged DNA is repaired before cells progress with the sporulation process or exit from spores (27,42). DisA is encoded in a multi-gene operon and the gene immediately upstream of *disA* codes for RadA (also referred to as SMS). *B. subtilis* RadA possesses 5’ to 3’ DNA helicase activity, contributes to DNA repair and DNA transformation processes in *B. subtilis* and has been shown to interact and negatively impact the activity of DisA (27). However, the mechanistic basis of how RadA binding to DisA inhibits the cyclase is currently not known. There is now ample evidence that the activity of the “house-keeping” membrane-linked c-di-AMP cyclase CdaA is impacted by two interacting proteins, the membrane-linked regulator protein CdaR and the cytoplasmic phosphoglucomutase enzyme GlmM (29,33). We have provided experimental evidence for the mechanistic basis by which GlmM inhibits the activity of CdaA, that is by preventing the formation of active higher-level oligomers. How CdaR regulates the cyclase activity of CdaA remains unclear. Recent work on the homologous proteins in *L. monocytogenes* indicated that the interaction of CdaA with CdaR takes place via the N-terminal transmembrane region of CdaA, GlmM has been shown to interact directly with the cytoplasmic cyclase domain of CdaA in *S. aureus* (28,33). Here, we show that this is also the case for the *B. subtilis* GlmM protein, which can bind without the requirement of any additional factor to the catalytic CdaA_CD_ domain. In future works, it will be interesting to determine the structure of the full-length CdaA enzyme, which might provide further insight into how the enzyme forms higher oligomers for activity as well as how it interacts with CdaR. Furthermore, it will be interesting to further investigate the interaction between GlmM and CdaR with CdaA within bacterial cells to determine if this interaction is dynamic and which stimuli will promote or prevent complex formation to fine-tune the synthesis of c-di-AMP. Identifying how interacting proteins regulate the activity of these important cyclases, will provide important insight how bacterial cells maintain proper levels of c-di-AMP under different growth conditions and in different environments.

### Experimental procedures

#### Bacterial strains and plasmid construction

All bacterial strains and primers used in this work are listed in Tables S3 and S4, respectively. pET28b-derived plasmids were constructed for the overproduction of the C-terminal catalytic domain of the *Bacillus subtilis* CdaA enzyme starting from amino acid Phe97 and referred to as CdaA_CD_, GlmM and the GlmM_F369_ variant comprising residues Met1 to Phe369 but lacking the C-terminal domain 4. To this end, the corresponding DNA fragments were amplified by PCR using *B. subtilis* strain 168 chromosomal DNA as template and primer pairs ANG2760/ANG2761 (*cdaA*_*CD*_), ANG2762/ANG2763 (*glmM*) and ANG2762/ANG2764 (*glmM*_*F369*_). The PCR products were purified, digested with NheI/BamHI (*cdaA*_*CD*_) or NcoI/XhoI (*glmM* and *glmM*_*F369*_) and ligated with pET28b, which had been cut with the same enzyme. CdaA_CD_ was cloned in frame with an N-terminal thrombin cleavable 6-histidine tag, while GlmM and GlmM_F369_ were cloned in frame with a C-terminal 6-histidine tag and a thrombin cleavage site was introduced in front of the His-tag as part of the primer sequence. The resulting plasmids pET28b-*his*-*cdaA*_*CD*_, pET28b-*glmM-his* and pET28b-*glmM*_*F369*_-*his* were initially recovered in *E. coli* XL1-Blue, yielding the strains ANG4583, ANG4584 and ANG4585 and subsequently transformed for protein expression into strain *E. coli* BL21(DE3), yielding strains ANG4597, ANG4598 and ANG4599, respectively. Plasmids pET28b-*his*-*cdaA_CD_*-*R126*, pET28b-*glmM-D194A-his,* pET28b-*glmM-D151A/E154A-his* were constructed for the expression of CdaA_CD_ and GlmM alanine substitution variants. The plasmids were constructed by QuikChange mutagenesis using pET28b-*his*-*cdaA*_*CD*_ and primer pair ANG3373/ANG3374 or plasmid pET28b-*glmM*-*his* and primer pairs ANG3381/ANG3382 and ANG3383/3384, respectively. The plasmids were initially recovered in *E. coli* XL1-Blue, yielding strains ANG5933, ANG5937, ANG5938 and subsequently introduced for protein expression into *E. coli* strain BL21(DE3) yielding strains ANG5940, ANG5944, ANG5945. In addition, plasmid pET28b-*glmM-D151A/E154A/D194A*-*his* for expression of a GlmM variant with a triple Asp151 (D151), *Glu154 (E154) and Asp195 (D195)* alanine substitution variant was constructed by QuikChange mutagenesis using plasmid pET28b-*glmM-D194A*-*his* as template and primer pair ANG3383/3384 to introduce the D151A and E154A mutations. Plasmid pET28b-*glmM-D151A/E154A/D194A*-*his* was recovered in *E. coli* strain XL1-Blue, yielding strain ANG5939 and subsequently introduced for protein expression into strain BL21(DE3) yielding strain ANG4946. The sequences of all plasmid inserts were verified by fluorescent automated sequencing at Eurofins.

### Protein expression, purification and quantification

Proteins CdaA_CD_, GlmM and GlmM_F369_ were expressed and purified from 1 L cultures as previously described (28). Briefly, when bacterial cultures reached an OD_600_ of approximately 0.6, protein expression was induced with 1 mM IPTG (final concentration) for 3 hours at 37°C. Cells were harvested by centrifugation, suspended in 20 mL of 50 mM Tris pH 7.5, 500 mM NaCl buffered and lysed using a French Press system. Lysates were clarified by centrifugation and the supernatant loaded onto a gravity flow column with 3 mL of Ni-NTA resin. Immobilized proteins were washed with 20 mL of 50 mM Tris pH 7.5, 500 mM NaCl, 50 mM imidazole buffer and eluted in 5 x 1 ml fractions using 50 mM Tris pH 7.5, 200 mM NaCl, 500 mM imidazole buffer. Fractions containing the proteins were pooled and loaded onto a preparative Superdex 200 HiLoad 16/60 column equilibrated with 1 column volume of 30 mM Tris pH 7.5, 150 mM NaCl buffer. When appropriate, the purified proteins were concentrated using 10 mL 10 kDa cutoff Amicon concentrators for downstream applications. For the purification of the CdaA_CD_:GlmM and CdaA_CD_:GlmM_F369_ complexes or complexes of CdaA_CD_ and GlmM alanine-substitution variants, cell lysates of strains overproducing the respective proteins were mixed after the French press step, then the same protein purification procedure steps as described above used for the purification of individual proteins were performed. Protein concentrations were determined using the BCA assay kit (Pierce™ BCA Protein Assay Kit). For each sample, the readings were taken in triplicates and then averaged to obtain the protein concentration. Purified proteins were also separated on 12% SDS PAGE gels and detected by Coomassie staining.

### Microscale Thermophoresis

A Microscale Thermophoresis (MST) experiment was performed to determine the binding affinity between the *B. subtilis* CdaA_CD_ and GlmM proteins. The CdaA_CD_ and GlmM were expressed and purified from 1 L cultures as described above, however using 20 mM HEPES, pH 7.5, 500 mM NaCl buffer for the Ni-NTA purification and 10 mM HEPES, pH 7.5, 150 mM NaCl for the SEC purification step. Next, CdaA_CD_ was fluorescently labelled with an amine-reactive dye using the Monolith Protein Labelling RED-NHS 2nd Generation kit (NanoTemper Technologies GmbH). To this end, 90 μl of a 40 μM CdaA_CD_ solution was mixed with 10 μl of a 400 μM dye solution in 10 mM HEPES, pH 7.5, 150 mM NaCl, 0.05% Tween-20 buffer and incubated for 30 min at room temperature in the dark. Unincorporated dye was subsequently removed from the labelled protein as described in the manufacturer’s instructions. Following the labelling reaction, the protein concentration was determined by nanodrop and using the Beer-Lambert equation and an extinction coefficient of 0.774 for CdaA_CD_. For the MST experiment, a 50 nM solution of the fluorescently labelled CdaA_CD_ protein was mixed at a 1:1 ratio with a solution of purified GlmM protein at a starting concentration of 1600 μM and ten 2-fold dilutions there of prepared in the purification buffer (10 mM HEPES, pH 7.5, 150 mM NaCl, 0.05% Tween-20). The samples were filled into individual premium capillaries and subsequently loaded in the capillary tray. Each MST run was performed on a Monolith NT.115 instrument at a light emitting diode (LED) power of 95% and microscale thermophoresis (MST) power of 80% with a duration of 30 seconds laser on time (NanoTemper Technologies GmbH) (43). The experiment was performed 5 times and average normalized fluorescence values and standard deviations determined and plotted. For the data analysis and K_d_ determination the NT Analysis Software (NanoTemper Technologies GmbH) was used (43).

### Protein crystallisation, data processing and analysis

For crystallisation, the histidine tag was removed from the purified *B. subtilis* CdaA_CD_ protein. This was done by incubating 10 mg purified protein with 20 U thrombin overnight at 4°C with agitation. The following day, the tag less CdaA_CD_ was purified by size-exclusion chromatography as described above. The CdaA_CD_ protein was crystallized at a concentration of 4 mg/ml in 0.1M sodium cacodylate pH 6.5, 0.1M ammonium sulfate, 0.3 M sodium formate, 6% PEG 8000, 3% γ-PGA via the vapour diffusion method. The crystal screens for *B. subtilis* GlmM (including the His tag) were set up at a concentration of 10 mg/ml and protein crystals were obtained in two different conditions. The structure with bound PO_4_ (GlmM:PO_4_) was obtained in the Morpheus screen containing 0.1 M buffer system 1 (Imidazole; MES, pH 6.5), 0.09 M NPS (NaN03; Na_2_HPO_4_; (NH_4_)_2_SO_4_) and 37.5% MPD_P1K_P3350 (75% MPD, PEG 1K, PEG 3350) and the divalent-ion bound crystal structure (GlmM:metal ion) was obtained in 0.05 M Magnesium chloride hexahydrate, 0.1 M HEPES pH 7.5, 30% v/v Polyethylene glycol monomethyl ether 550 buffer. The crystals for the CdaA_CD_:GlmM_F369_ complex were set up at a protein concentration of 10 mg/ml and crystals were obtained in 0.12 M alcohols, 0.1 M buffer system (Imidazole; MES, pH 6.5) and 30% GOL_P4K (60% glycerol, PEG 4K) (Complex 1) and 0.1 M carboxylic acids, 0.1 M buffer system 1 (Imidazole; MES, pH 6.5) and 30% GOL_P4K (60% glycerol, PEG 4K) (Complex 2). The crystals were fished and stored in liquid nitrogen to test for diffraction at the I03 beamline at the Diamond Light Source (Harwell Campus, Didcot, UK). Data were reduced with DIALS (44) and scaled and merged with AIMLESS (45). The structures of CdaA_CD_ and GlmM were solved by the molecular replacement method using the program MR-PHASER (46) in the Phenix suite (47), using the *L. monocytogenes* CdaA structure (PDB 4RV7; (21)) and *B. anthracis* GlmM structure (PDB 3PDK; (37)) as the search models, respectively. To solve the phase problem for the structure of the CdaA_CD_:GlmM_F369_ complex, dimers of *B. subtilis* CdaA_CD_ and GlmM (each) were used as the search models using the MR-PHASER program in Phenix. The models were manually built using COOT (48) followed by iterative cycles of structure refinement using the Phenix-Refine program (49). The final refined structures were analysed using the PDBePISA server (36) to identify buried interface areas for each protein. To search for conserved residues among the phylogenetically related homologs, a protein BLAST search was performed using *B. subtilis* CdaA_CD_ and GlmM amino acid sequences as query sequences and a multiple sequence alignment (MSA) of the top 2500 homologs found in Firmicutes was prepared. The MSA was then used to identify conserved residues among the homologs using the ConSurf server (50).

### CdaA_CD_ activity and inhibition assays

To assess the activity of the *B. subtilis* CdaA_CD_ enzymes, 20 μl enzyme reactions were set up in 100 mM NaCl, 40 mM HEPES pH 7 buffer containing 10 mM MnCl_2_ (or 10 mM MgCl_2_ or 10 mM CoCl_2_ for metal dependent assays), 100 mM ATP spiked with a-P^32^-labelled ATP (Perkin Elmer; using 0.4 μl of a 3.3 μM, 250 μCi solution per 20 μl reaction) and 5 μM CdaA_CD_ enzyme. The mixture was incubated at 37°C for 4 hours, followed by heat inactivation at 95°C for 5 minutes. After centrifugation for 10 minutes at 21,000 x g, 1 μl of the mixture was deposited onto a polyethylenimine-modified cellulose TLC plate (Millipore) and nucleotides separated by running the plate for 20 minutes using a 3.52 M (NH_4_)_2_SO_4_ and 1.5 M KH_2_PO_4_ buffer system mixed at a 1:1.5 v/v ratio. Radioactive signals for ATP and the c-di-AMP reaction product were detected using a Typhoon FLA-700 phosphor imager. The bands were quantified using the ImageQuant program and the obtained values used to calculate the percent conversion of ATP to c-di-AMP. For the time course experiment, a 100 μl reaction mixture was prepared as described above and incubated at 37°C. Ten μl aliquots were removed at the indicated time points and the enzyme reactions stopped by incubation the removed aliquots at 95°C for 5 minutes. To assess the activity of CdaA_CD_ in the presence of GlmM or GlmM_F369_, the full length GlmM protein or C-terminally truncated GlmM variant were added to the reaction mixture at a 1:2 (CdaA_CD_: GlmM or CdaA_CD_: GlmM_F369_) molar ratio and the reactions incubated at 37°C for 4 hours, stopped and analysed as described above. The enzyme activity assays were performed in triplicates with two independently purified protein preparations.

### SAXS sample preparation and analysis

For the SAXS analysis, purified CdaA_CD_, GlmM, CdaA_CD_:GlmM complex and CdaA_CD_:GlmM_F369_ complex protein samples where purified by size exclusion chromatography as described above and subsequently concentrated to 5 mg/ml for CdaA_CD_, 24 mg/ml each for GlmM and the CdaA_CD_:GlmM complex and 10 mg/ml for the CdaA_CD_:GlmM_F369_ complex. Next, 50 μl protein samples were loaded on a high pressure Shodex column (KW403: range 10 kDa to 700 kDa) fitted to an Agilent 1200 HPLC system at the B21 beamline at the Diamond Light Source (Didcot, UK). The size-exclusion column was equilibrated with 30 mM Tris pH 7.5, 150 mM NaCl buffer prior to loading the protein sample and the data were collected continuously throughout the protein elution. The analysis of the datasets was done via ScÅtter (51) using the scattering frames corresponding to the elution peaks. The ab-initio analysis of the SAXS data to reconstruct a low-resolution shape of the model was done using DAMAVER (DAMMIF) program (52) which performs 13 *ab-initio* runs to generate models from each run that were averaged to determine the most persistent three-dimensional shape of the protein. The cross-correlation Normalised Spatial Discrepancy (NSD) values were calculated using DAMAVER (DAMSEL)(52)from each of the 13 generated models. The mean NSD values calculated for each of the protein were: 0.591 ± 0.088 (CdaA_CD_), 0.598 ± 0.014 (GlmM), 1.201 ± 0.099 (CdaA_CD_:GlmM_F369_ complex) and 0.661 ± 0.064 (CdaA_CD_:GlmM complex). The program Chimera (53) was used to visualise the reconstructed SAXS maps. The crystal structures of CdaA_CD_ and GlmM dimers as well as CdaA_CD_:GlmM_F369_ complex were fitted in the respective SAXS envelopes in Chimera using the one-step fit function. For the CdaA_CD_: GlmM complex data, a structural model of the CdaA_CD_ full-length GlmM complex was generated based on the crystal structure of CdaA_CD_: GlmM_F369_ complex, which was then fitted into the SAXS envelope data using Chimera.

## Data availability

The structure coordinates of the *B. subtilis* proteins were deposited in the Protein Database (https://www.rcsb.org), under PDB codes 6HUW (CdaA_CD_), 7OJR (GlmM: PO_4_ bound), 7OML (GlmM: metal bound) and 7OLH (CdaA_CD_:GlmM_F369_ Complex 1) and 7OJS (CdaA_CD_:GlmM_F369_ Complex 2). SAXS models were deposited in the SASBDB database, under the accession codes SASDL25 (Complex CdaA_CD_:GlmM), SASDMQ5 (Complex CdaA_CD_:GlmM_F369_), SASDLZ4 (GlmM) and SASDLY4 (CdaA_CD_).

## Supporting Information

This article contains a supporting information file with the following additional references (36,50,51,54,55).

## Acknowledgments

We would like to thank Christiaan van Ooij for helpful comments on the manuscript.

## Author contribution statement

**Monisha Pathania:** Conceptualization, Investigation, Data analysis, Visualization, Writing – original draft preparation. **Tommaso Tosi:** Conceptualization, Investigation, Data analysis, Writing – review & editing. **Charlotte Millsership:** Investigation, Data analysis, Writing – review & editing. **Fumiya Hoshiga:** Investigation, Data analysis, Writing – review & editing. **Rhodri M. L. Morgan:** Data analysis, Supervision, Writing – review & editing. **Paul S. Freemont:** Conceptualization, Funding acquisition, Supervision, Data analysis, Writing – review & editing. **Angelika Gründling:** Conceptualization, Funding acquisition, Data analysis, Supervision, Writing – original draft preparation.

## Funding and additional information

This work was funded by the MRC grant MR/P011071/1 to AG and PSF and the Wellcome Trust grant 210671/Z/18/Z to AG. The crystallization facility at Imperial College was funded by the BBSRC (BB/D524840/1) and the Wellcome Trust (202926/Z/16/Z). X-Ray datasets for CdaA_CD_, GlmM and the CdaA_CD_:GlmM complex 2 were collected at the I03 beamline and the CdaA_CD_:GlmM complex 1 dataset was collected at the I04 beamline at the Diamond Light Source (Didcot, UK). The SAXS data were collected at the B21 beamline at the Diamond Light Source (Didcot, UK).

## Conflict of Interest

The authors declare no conflicts of interest in regard to this manuscript.

## Notes

### Competing Interest Statement

The authors have declared no competing interest.

### Summary of Updates

Additional experiments were performed to determine the binding affinity between GlmM and CdaACD and an additional SAXS experiment was performed. The Discussion section was expanded.

