## Supporting Information for "Structural basis for the inhibition of the *Bacillus subtilis* c-di-AMP cyclase CdaA by the phosphoglucomutase GlmM"

List of Material included:
Supplemental Table 1: Interactions between $CdaA_{CD}$ and $GlmM_{F369}$ in the complex 1 structure as assessed with PDBePISA
Supplemental Table 2: SAXS data statistics
Supplemental Table 3: Bacterial strains used in this study
Supplemental Table 4: Primers used in this study
Supplemental Figure S1: Overlap of *B. subtilis, L. monocytogenes* and *S. aureus* CdaA/DacA catalytic domain structures.
Supplemental Figure S2: *B. subtilis* GlmM structure.
Supplemental Figure S3: Crystal structure of the *B. subtilis* $CdaA_{CD}$:$GlmM_{F369}$ complex 2.
Supplemental Figure S4: Overlay of $CdaA_{CD}$:$GlmM_{F369}$ complex structure.
Supplemental Figure S5: Amino acid residues fomring interactions between $CdaA_{CD}$ and $GlmM_{F369}$ in complex 1.
Supplemental Figure S6: SAXS scatter curves and SEC-SAXS elution profiles of the purified $CdaA_{CD}$, GlmM and $CdaA_{CD}$:GlmM proteins.
Supplemental Figure S7: Effect of amino acid substitutions in GlmM on *in vitro* complex formation with $CdaA_{CD}$.

**Supplemental Tables**

**Table S1:** Interactions between $CdaA_{CD}$ and $GlmM_{F369}$ in the complex 1 structure as assessed with PDBePISA (1)

| **$CdaA_{CD}$ Monomer 1 Residue (Atom)** | **$GlmMF_{369}$ Monomer 1 Residue (Atom)** | **Distance in Å** | **Type of interaction** |
|---|---|---|---|
| Tyr 122 (OH) | Glu 154 (OE1) | 2.90 | H-bond |
| Arg126 (NH2) | Glu 154 (OE2) | 3.91 | Ionic bond |
| Arg126 (NE) | Glu 154 (OE1) | 3.26 | Ionic bond |
| Arg 126 (NH2) | Asp 151 (O) | 2.69 | Ionic bond |
| Arg 126 (NH1) | Asp 151 (O) | 2.80 | Ionic bond |
| Arg127 (O) | Lys158 (NZ) | 3.75 | H-bond |
| Asn166 (OD1) | Gln 161 (N) | 3.72 | H-bond |
| Asn166 (ND2) | Gln 157 (OE1) | 2.96 | H-bond |
| **$CdaA_{CD}$ Monomer 2 Residue (Atom)** | **$GlmMF_{369}$ Monomer 1 Residue (Atom)** | **Distance in Å** | **Type of interaction** |
| Arg127 (NH1) | Ser186 (O) | 2.92 | H-bond |
| Arg127 (NE) | Thr190 (OG1) | 3.23 | H-bond |
| Arg127 (NH1) | Thr190 (OG1) | 3.00 | H-bond |
| Arg126 (O) | Thr190 (OG1) | 3.86 | H-bond |
| Arg126 (O) | His 191 (ND1) | 2.81 | H-bond |
| Arg126 (NE) | Asp195 (OD1) | 3.09 | Ionic bond |
| Arg126 (NH1) | Asp195 (OD1) | 2.63 | Ionic bond |
| Arg126 (NH2) | Asp195 (OD1) | 3.78 | Ionic bond |
| **$CdaA_{CD}$ Monomer 2 Residue (Atom)** | **$GlmMF_{369}$ Monomer 2 Residue (Atom)** | **Distance in Å** | **Type of interaction** |
| Glu145 (OE1) | Asp195 (OD1) | 3.44 | H-bond |
| Glu145 (OE2) | Asp195 (OD1) | 3.10 | H-bond |

For amino acid atom labelling shown in brackets, see Figure S5.

**Table S2:** SAXS data statistics

| | **$CdaA_{CD}$** | **GlmM** | **$CdaA_{CD}$/GlmM Complex** | **$CdaA_{CD}$/$GlmM_{F369}$ Complex** |
|---|---|---|---|---|
| Beamline | B21 | B21 | B21 | B21 |
| Wavelength (Å) | 1 | 1 | 1 | 1 |
| q-range ($Å^{-1}$) | 0.0032-0.38 | 0.0032-0.38 | 0.0032-0.38 | 0.0032-0.38 |
| I(0) ($cm^{-1}$) | $0.0027 \pm 1.6 \text{ x}10^{-5}$ | $0.25 \pm 9.7 \text{ x}10^{-5}$ | $0.53 \pm 1.9 \text{ x}10^{-4}$ | $0.11 \pm 1.2 \text{ x}10^{-4}$ |
| Rg (Å) | 26.28 ± 0.19 | 38.21 ± 0.02 | 44.75 ± 0.02 | 37.39 ± 0.22 |
| Dmax (Å) | 88 | 122 | 165 | 117.5 |
| Porod Volume (Å3) | 112507 | 256783 | 394542 | 215271 |
| MW Estimate (kDa) | 42 | 85 | 130 | 97.8 |
| No. of atoms (dammif) | 1931 | 1900 | 1855 | 2043 |

**Table S3:** Bacterial strains used in this study

| **Unique ID** | **Relevant features** | **Reference** |
|---|---|---|
| ***Escherichia coli* strains** | | |
| ANG127 | XL1-Blue - Cloning strain; TetR | Stratagene |
| ANG191 | BL21(DE3) - Protein expression strain | Novagen |
| ANG4583 | XL1-Blue pET28b-*his-cdaA*$_{CD}$; KanR | This study |
| ANG4597 | BL21(DE3) pET28b-*his-cdaA*$_{CD}$; KanR | This study |
| ANG4584 | XL1-Blue pET28b-*glmM-his*; KanR | This study |
| ANG4598 | BL21 (DE3) pET28b-*glmM-his*; KanR | This study |
| ANG4585 | XL1-Blue pET28b-*glmMF369-his*; KanR | This study |
| ANG4599 | BL21(DE3) pET28b-*glmMF369-his*; KanR | This study |
| ANG5933 | XL1-Blue pET28b-*his-cdaA*$_{CD}$*-R126A*; KanR | This study |
| ANG5937 | XL1-Blue pET28b-*glmM-D195A-his*; KanR | This study |
| ANG5938 | XL1-Blue pET28b-*glmM-D151A/E154A-his*; KanR | This study |
| ANG5939 | XL1-Blue pET28b-*glmM-D151A/E154A/D195A-his*; KanR | This study |
| ANG5940 | BL21 (DE3) pET28b-*his-cdaA*$_{CD}$*-R126A*; KanR | This study |
| ANG5944 | BL21 (DE3) pET28b-*glmM-D195A-his*; KanR | This study |
| ANG5945 | BL21 (DE3) pET28b-*glmM-D151A/E154A-his*; KanR | This study |
| ANG5946 | BL21 (DE3) pET28b-*glmM-D151A/E154A/D195A-his*; KanR | This study |
| ***Bacillus subtilis strains*** | | |
| ANG196 | *B. subtilis* strain 168 | (2) |

Antibiotics were used at the following concentrations: Kanamycin (KanR) 30 µg/ml and 5-10 µg/ml tetracycline (TetR)

**Table S4:** Primers used in this study

| Number | Name | Sequence |
|---|---|---|
| ANG2760 | 5-NheI-BSCdaA-97 | CTAGCTAGCTTTTTTTCGAGGAGCGGCACGCCTG |
| ANG2761 | 3-BamHI -BScdaA-stop | CGGGATCCTTATCCATTTTTCTTGCCCCTCCAATACC |
| ANG2762 | 5-NcoI-BSGlmM | CACGCCATGGGCAAGTATTTTGGAACAGACGGTGT AAG |
| ANG2763 | 3-XhoI-BSGlmM | CCGCTCGAGGCCGCTGCTGCCGCGCGGCACCAGCTC TAATCCCATTTCTGACCGGACGAC |
| ANG2764 | 3-XhoI-BSGlmMF369 | CCGCTCGAGGCCGCTGCTGCCGCGCGGCACCAGCTC TAATCCCATTTCTGACCGGACGAC |
| ANG3373 | 5-CdaA(BS)_R126A | GCAATCAATTATATGGCGAAAGCCCGTATAGGCGC CCTGCTGAC |
| ANG3374 | 3-CdaA(BS)_R126A | GTCAGCAGGGCGCCTATACGGGCTTTCGCCATATAA TTGATTGC |
| ANG3381 | 5-GlmM(BS)_D195A | GCGACACACCTGTTTGCTGCTTTAGATGCAGATGTT TCTAC |
| ANG3382 | 3-GlmM(BS)_D195A | GTAGAAACATCTGCATCTAAAGCAGCAAACAGGTG TGTCGC |
| ANG3383 | 5-GlmM(BS)_D151A-E154A | CAGACCTTGGACTTGTAAACGCTTATTTTGCAGGCG GACAAAAATATCTGCAATTC |
| ANG3342 | 3-GlmM(BS)_D151A-E154A | GAATTGCAGATATTTTTGTCCGCCTGCAAAATAAGC GTTTACAAGTCCAAGGTCTG |

Restriction sites in primer sequences are underlined.

**Supplemental figures and legends**

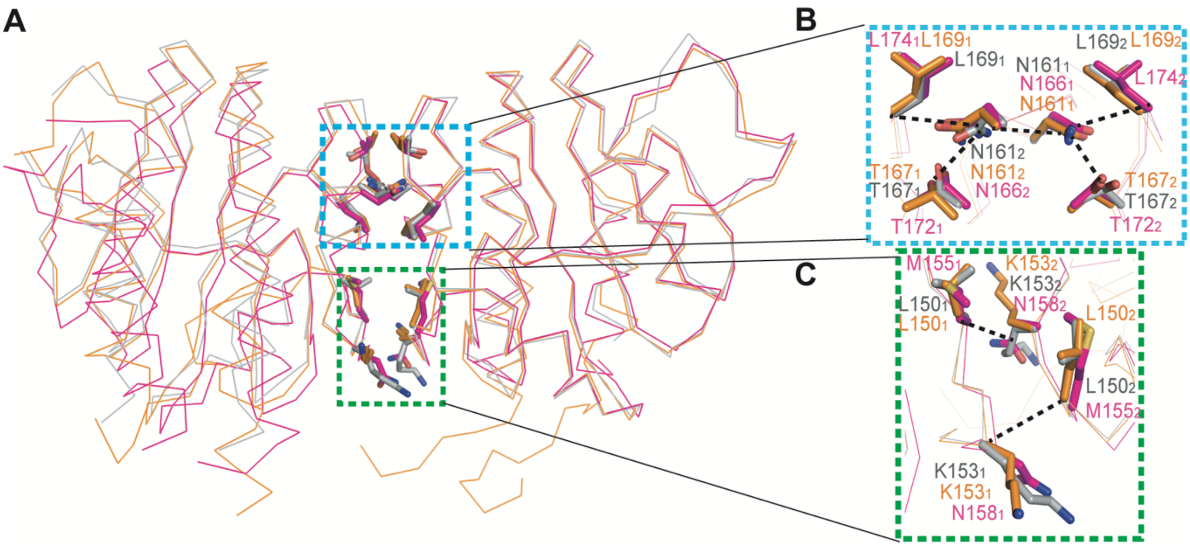

**Figure S1: Overlap of *B. subtilis, L. monocytogenes* and *S. aureus* CdaA/DacA catalytic domain structures.** *A,* Superimposition of *B. subtilis* CdaA$_{CD}$ (grey), *L. monocytogenes* Δ100CdaA$_{CD}$ (yellow; PDB 4RV7) and *S. aureus* DacA$_{CD}$ (pink; PDB 6GYW) structures in line representation with dimer interface residues shown in stick mode. *B* and *C*, zoomed in views of the interacting residues at the dimer interface.

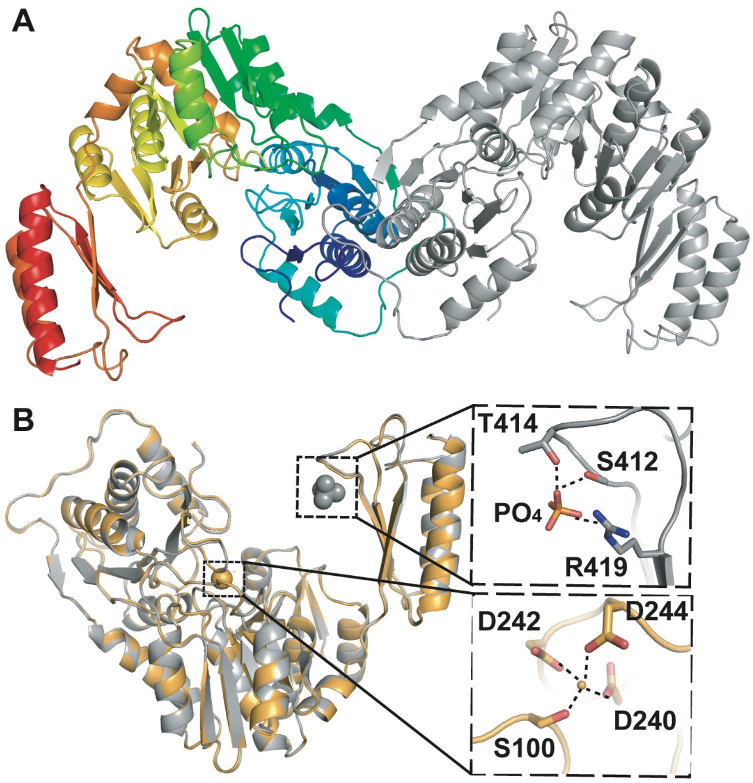

**Figure S2. *B. subtilis* GlmM structure.** *A,* Cartoon representation of the *B. subtilis* GlmM dimer with one monomer shown in rainbow colours and the other one in grey. *B,* Overlay of the *B. subtilis* GlmM metal-bound (shown in gold) and phosphate-bound (shown in grey) structures.

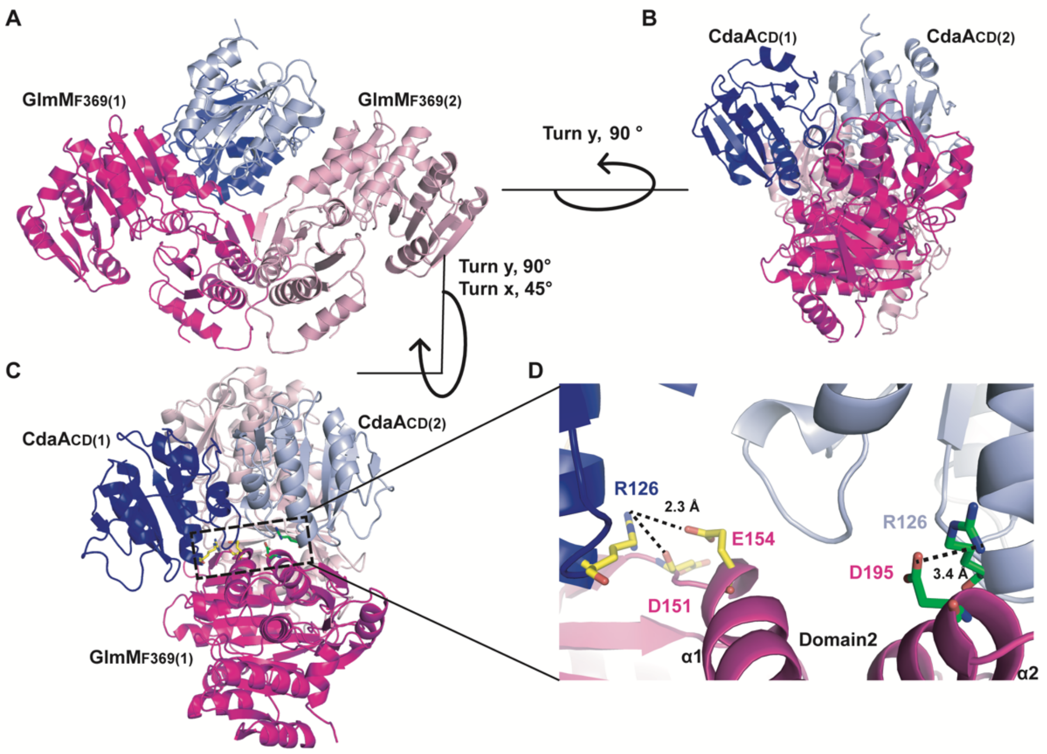

**Figure S3: Crystal structure of the *B. subtilis* $CdaA_{CD}$:$GlmM_{F369}$ complex 2.** (*A-D*) Structure of the *B. subtilis* $CdaA_{CD}$:$GlmM_{F369}$ lower resolution complex 2 shown in cartoon representation. The *B. subtilis* $CdaA_{CD}$:$GlmM_{F369}$ complex crystallized as a dimer of dimers with individual $GlmM_{F369}$ monomers shown in dark pink [$GlmM_{F369}$(1)] and light pink [$GlmM_{F369}$(2)] and individual $CdaA_{CD}$ monomers shown in dark blue [$CdaA_{CD}$(1)] and light blue [$CdaA_{CD}$(2)], respectively. The complex is show in *A,* in front view, *B,* in side view (rotated 90º along the y-axis) and *C*, in top-side view rotated at the angle as indicated with respect to panel A. *D,* Zoomed in view of the $CdaA_{CD}$/$GlmM_{F369}$ interface. Residue Arg126 from $CdaA_{CD}$(1) forms H-bond and ionic interactions with Asp151 and Glu154 of $GlmM_{F369}$(1) (residues shown in yellow), and residue Arg126 from $CdaA_{CD}$(2) forms ionic interactions with Asp 195 in $GlmM_{F369}$(1) (residues shown in green). The images were prepared in PyMOL (The PyMOL Molecular Graphics System, Version 2.0 Schrödinger, LLC).

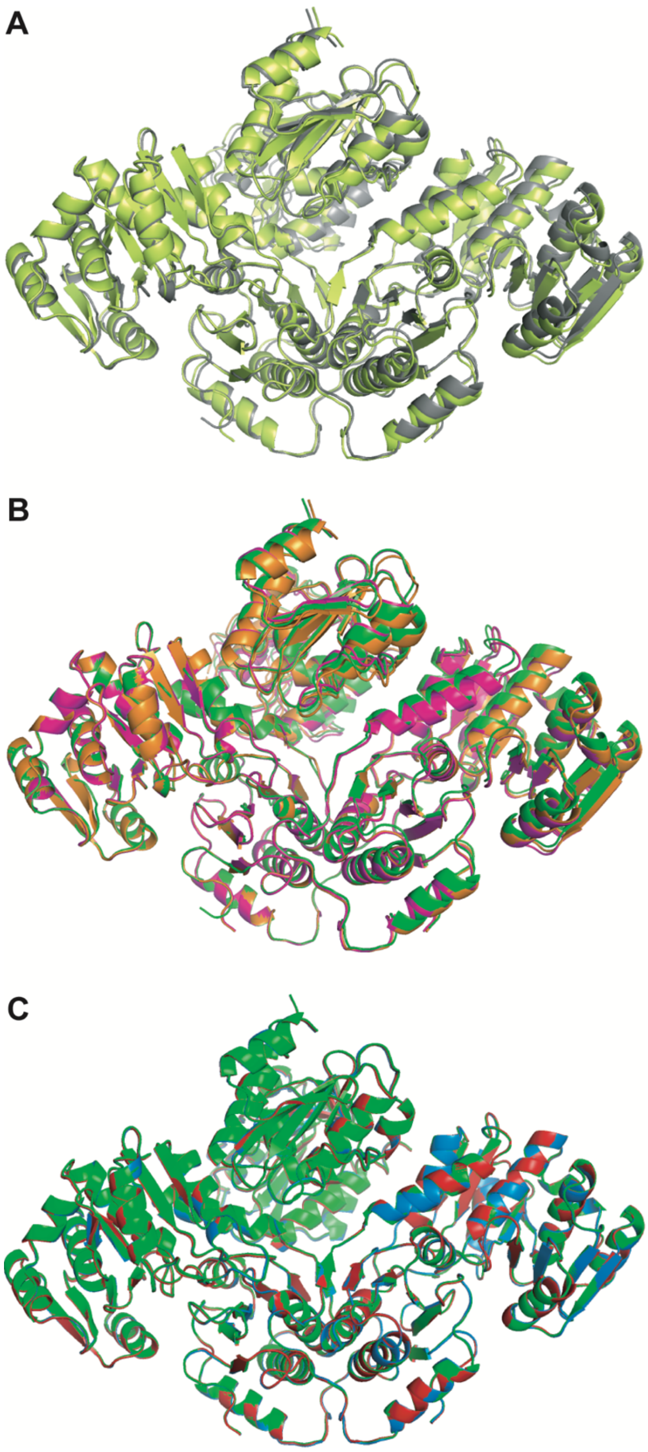

**Figure S4: Overlay of CdaA$_{CD}$:GlmM$_{F369}$ complex structure.** *A)* Overlay of the 3.6 Å resolution CdaA$_{CD}$:GlmM$_{F369}$ complex 1 (PDB 7OLH; grey) and the 4.2 Å resolution CdaA$_{CD}$:GlmM$_{F369}$ complex 2 (PDB 7OJS; light green) structures. *B)* Overlay of the three CdaA$_{CD}$/GlmM$_{F369}$ complexes present within the asymmetric unit of the complex structure 1 (PDB 7OLH) with complexes shown in pink, orange and green colors. The complexes overlayed with r.m.s.d. scores of 0.15 and 0.20 Å. *C)* Overlay of the three CdaA$_{CD}$/GlmM$_{F369}$ complexes present within the asymmetric unit of the complex structure 2 (PDB 7OJS) with complexes shown in red, green and blue, respectively. The complexes overlayed with r.m.s.d. scores of 0.15-0.16 Å.

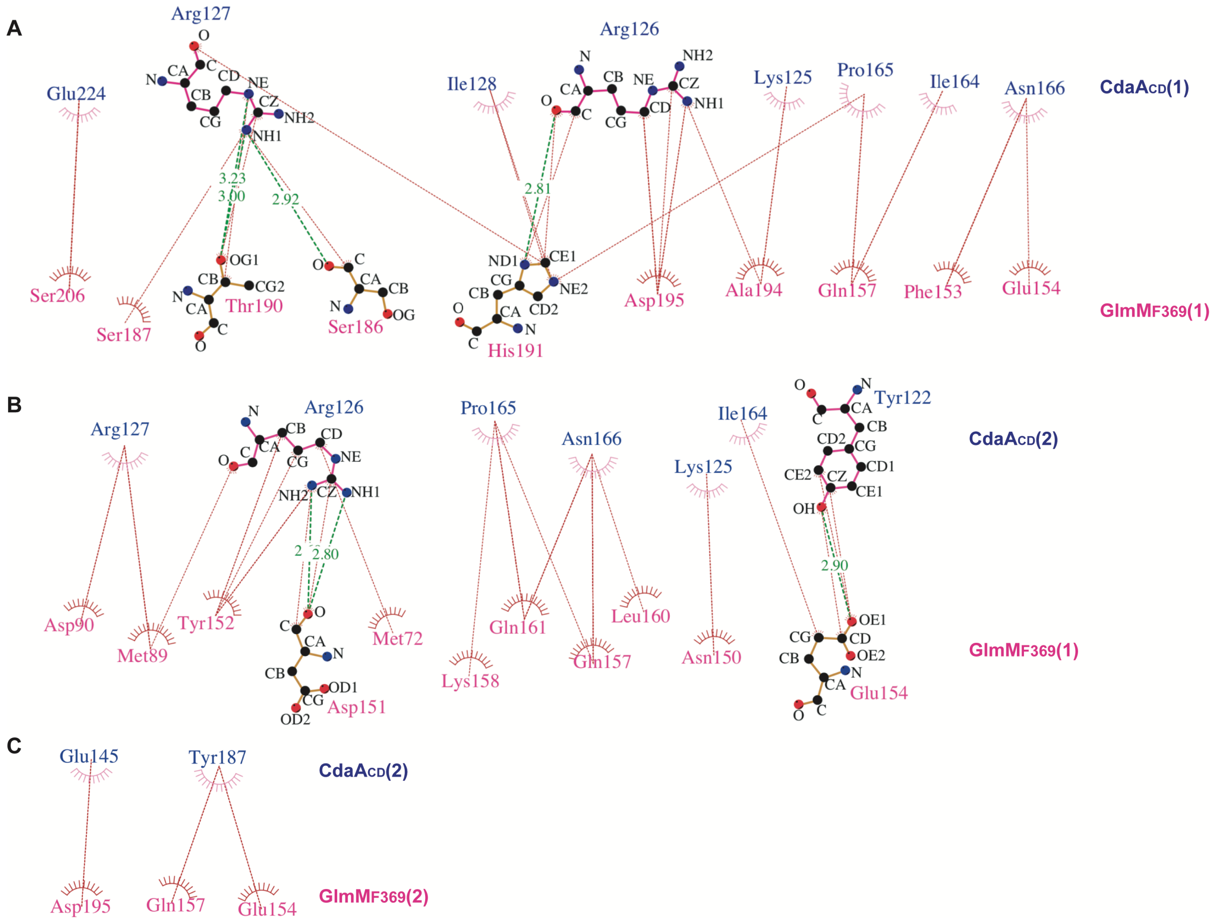

**Figure S5: Amino acid residues fomring interactions between CdaA$_{CD}$ and GlmM$_{F369}$ in complex 1.** (*A-C*) Schematic 2D-representation of amino acids interacting in *A,* CdaA$_{CD}$(1) with GlmM$_{F369}$(1), *B,* CdaA$_{CD}$(2) with GlmM$_{F369}$(1) and *C,* CdaA$_{CD}$(2) with GlmM$_{F369}$(2). The plots were prepared using LigPlot+ (3). The hydrogen bonds are displayed in green dashed lines, along with the distance shown in Å and hydrophobic bonds are represented in red dashed lines. The residues showed with the amino acid structure form a hydrogen bonding network as a part of the complex interface.

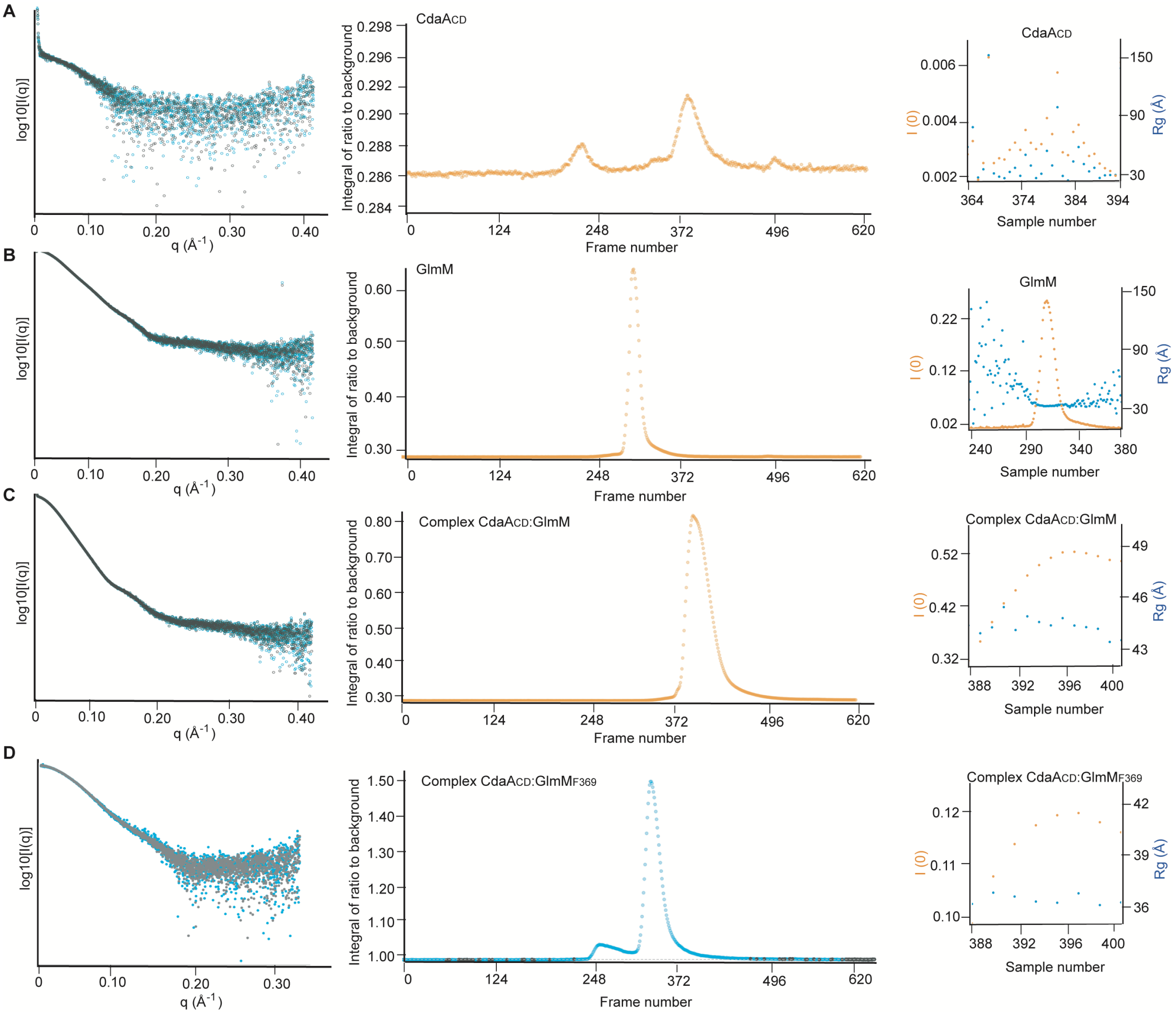

**Figure S6: SAXS scatter curves and SEC-SAXS elution profiles of the purified CdaA$_{CD}$, GlmM, CdaA$_{CD}$:GlmM complex and CdaA$_{CD}$:GlmM$_{F369}$ complex.** (*A-D*), 50 µl of *A,* CdaA$_{CD}$, *B,* GlmM, *C,* the CdaA$_{CD}$:GlmM complex or *D*, the CdaA$_{CD}$:GlmM$_{F369}$ complex were injected onto a Shodex column (KW403: range 10 kDa to 700 kDa) fitted to an Agilent 1200 HPLC system at the B21 beamline at the Diamond Light Source (Didcot, UK). *Left panels*: Log 10 intensity plot of subtracted and merged SAXS frames are shown, with black dots representing the averaged buffer frames subtracted from the averaged sample frames and cyan representing the median of the buffer frames subtracted from the averaged sample frames. *Middle panels*: A full dataset of 620 scattering frames was collected and the data were analyzed with ScÅtter (4) and scattering curves calculated. *Right panels,* double Y plot with zero-angle scattering intensity [I(0)] in orange, and radius of gyration (Rg) in cyan, estimated from the Guinier region for each subtracted frame using the program ScÅtter. Scattering frames were selected according to homogeneity of the estimated Rg values.

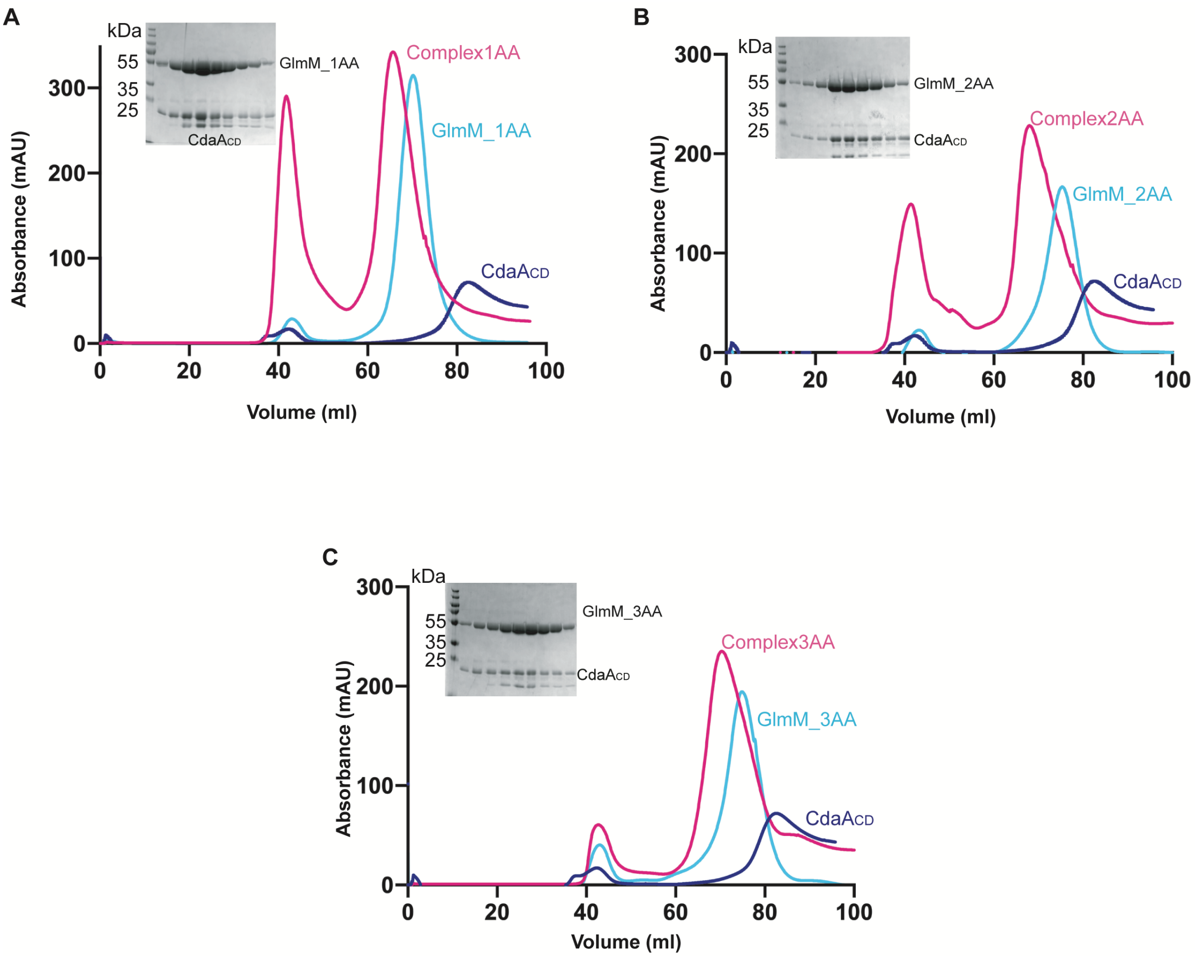

**Figure S7: Effect of amino acid substitutions in GlmM on *in vitro* complex formation with CdaA$_{CD}$.** (*A*-*C*), FPLC chromatograms and SDS-PAGE gel analysis of the *B. subtilis* complex formed between CdaA$_{CD}$ and the GlmM variant *A,* GlmM-D195A (GlmM_1AA), *B,* GlmM-D151A/E154A (GlmM_2AA) and *C*, GlmM-D151A/E154A/D195A (GlmM_3AA). The FPLC elution profiles were recorded at a wavelength of 280 nm and are shown for CdaA$_{CD}$ in blue, the GlmM variant in cyan and the complex in pink. Proteins from the peak fractions of each complex were separated on a 12% SDS PAGE gel and proteins visualized by Coomassie staining (shown in the inserts). The experiment in panels A and B was performed once and for panel C one representative result from two experiments is shown.

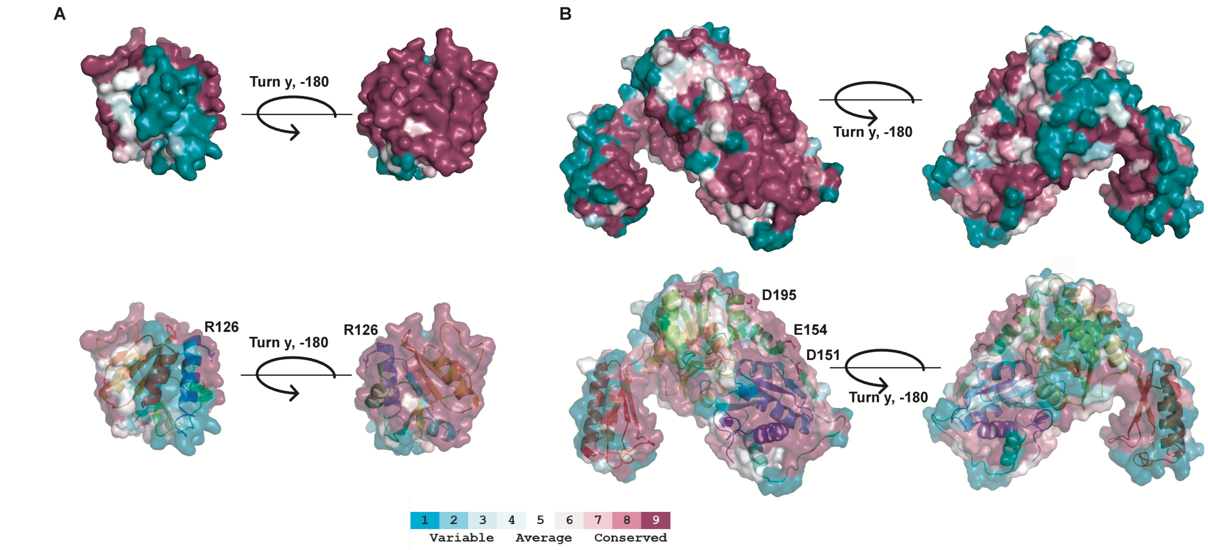

**Figure S8: Consurf models of CdaA and GlmM.** *A)* A protein BLAST search was performed using the *B. subtilis* $CdaA_{CD}$ amino acid sequence as query sequence and a multiple sequence alignment (MSA) of the top 250 homologs found in Firmicutes prepared. The MSA was then used to identify conserved residues among the homologs using the ConSurf server (5). In the top panel, the Consurf-surface view of $CdaA_{CD}$ is shown in two different orientations rotated by 180 degree along the Y-axes. In the bottom panel, a combined Consurf-surface view and cartoon representation of $CdaA_{CD}$ in rainbow colours is shown with residue R126 shown in stick mode. *B*) Same as in panel A but using the *B. subtilis* GlmM amino acid sequence as query sequence. In the bottom panel, GlmM residues D151, E154 and D195 are shown in stick mode. Highly conserved residues are shown in dark purple and least conserved residues in turquoise as indicated in the bar shown below the panels.
